# The low complexity regions in the C-terminus are essential for the subcellular localisation of *Leishmania* casein kinase 1 but not for its activity

**DOI:** 10.1101/2020.02.28.969741

**Authors:** Daniel Martel, Stewart Pine, Katharina Bartsch, Joachim Clos, Gerald F. Späth, Najma Rachidi

## Abstract

Casein Kinase 1 (CK1) family members are serine/threonine protein kinases ubiquitously expressed in eukaryotic organisms. They are involved in a wide range of important cellular processes, such as membrane trafficking, or vesicular transport in organisms from yeast to humans. Due to its broad spectrum of action, CK1 activity and expression is tightly regulated by a number of mechanisms, including subcellular sequestration. Defects in CK1 regulation, localisation or the introduction of mutations in the CK1 coding sequence are often associated with important diseases such as cancer. Increasing evidence suggest that the manipulation of host cell CK1 signalling pathways by intracellular pathogens, either by exploiting the host CK1 or by exporting the CK1 of the pathogen into the host cell may play an important role in infectious diseases. *Leishmania* CK1.2 is essential for parasite survival and released into the host cell, playing an important role in host pathogen interactions. Although *Leishmania* CK1.2 has dual role in the parasite and in the host cell, nothing is known about its parasitic localisation and organelle-specific functions. In this study, we show that CK1.2 is a ubiquitous kinase, which is present in the cytoplasm, associated to the cytoskeleton and localised to various organelles, indicating potential roles in kinetoplast and nuclear segregation, as well as ribosomal processing and motility. Furthermore, using truncated mutants, we show for the first time that the two low complexity regions (LCR) present in the C-terminus of CK1.2 are essential for the subcellular localisation of CK1.2 but not for its kinase activity, whereas the deletion of the N-terminus leads to a dramatic decrease in CK1.2 abundance. In conclusion, our data on the localisation and regulation of *Leishmania* CK1.2 contribute to increase the knowledge on this essential kinase and get insights into its role in the parasite.

## Introduction

Casein Kinase 1 (CK1) family members are serine/threonine protein kinases ubiquitously expressed in eukaryotic organisms [1]. They are involved in a wide range of important cellular processes, such as membrane trafficking, or vesicular transport in organisms from yeast to humans [1]. Due to its broad spectrum of action, CK1 activity and expression are tightly regulated by a number of mechanisms, including subcellular sequestration or phosphorylation [1]. Defects in CK1 regulation, localisation or the introduction of mutations in the CK1 coding sequence are often associated with important diseases such as cancer [2] [3] [4]. Consistent with its various roles, members of the CK1 family are associated with many subcellular structures. In mammalian cells, CK1δ has been detected at the centrosomes and the *trans*-Golgi network, performing an important role as a mediator of ciliogenesis [5, 6]. CK1δ as well as CK1α interact with membrane structures of the endoplasmic reticulum, the Golgi and various transport vesicles [5] [7]. In budding yeast, ScHrr25/CK1δ is localised to the bud neck, where it is essential for proper cytokinesis, and to endocytic sites, where it is required for their initiation and stabilisation [8] [9] [10]. Lastly, ScHrr25/CK1δ is recruited to cytoplasmic processing bodies (P-bodies), which protects the active kinase from the cytoplasmic degradation machinery during stress [11]. These few examples reflect the tight association of CK1 localisation to its functions, suggesting that investigating its localisation may increase our knowledge on this kinase and allow the identification of potential novel functions.

Increasing evidence suggests that the manipulation of host cell CK1 signalling pathways by intracellular pathogens, either by exploiting host CK1 or by exporting the CK1 of the pathogen into the host cell might play an important role in infectious diseases [12] [13] [14] [15] [16]. Indeed, host CK1 pathways are vital for *Mycobacterium* [15]. Knockdown of host CK1 leads to the decrease of infectious bursal disease virus (IBDV) replication [17]. The relationship between CK1 and viral replication was also demonstrated for other viruses, such as Simian Virus 40, hepatitis C virus and yellow fever virus [18, 19], [20]. *Plasmodium falciparum* CK1 is essential for parasite survival as well as released into the host cell [21]. Indeed, PfCK1 has a role in invasion through its interaction with and phosphorylation of PfRON3 (RhOptry Neck protein 3), which is located in the rhoptries [12]. *Leishmania* CK1.2 is also essential for parasite survival and was identified in exosomes [13] [22]. Thus, *Leishmania*, as the causative agent of Leishmaniasis, represents an excellent model to study the cellular roles of CK1. This parasite has two developmental stages, an extracellular promastigote that proliferated inside the insect vector, and an intracellular amastigote that develops and multiply inside the phagolysosomes of macrophages. There are six members of the CK1 family in *Leishmania* and little is known about their localisation or functions as only two paralogs were studied: CK1.4 and CK1.2. CK1.4 is mainly localised in the cytoplasm, and unlike other *Leishmania* CK1s contains a putative secretion signal. This paralog, secreted by the parasite was shown to be important for virulence [23]. CK1.2 is the major CK1 paralog, the most conserved kinase in *Leishmania spp.* and the most closely related to its human orthologs, suggesting that it might have been selected by the host cell rather than by the parasite. Lastly, *Leishmania* CK1.2, released in the host cell as free protein or via exosomes was shown to phosphorylate the host IFNAR1 (a receptor for alpha/beta interferon), leading to its degradation and the attenuation of the cellular responses to IFN-α, mimicking human CK1α [22, 24, 25] [26]. These data suggest that *Leishmania* CK1.2 phosphorylates host proteins to subvert macrophages and favours parasite survival, making this kinase a key player in host-pathogen interactions. Despite its essential role in parasite viability and virulence through its dual role in parasite and host cell, nothing is known about the functions of CK1.2 in the parasite [13] [25].

In this study, we show that CK1.2 is a ubiquitous protein kinase, present in the cytoplasm and associated to the cytoskeleton. In addition, CK1.2 localises to various organelles, such as the basal body, the flagellum, and the nucleolus, suggesting potential functions in the regulation of kinetoplast and nuclear segregation and/or ribosomal processing. Furthermore, using truncated mutants, we show for the first time that the two low complexity regions (LCR) present at the C-terminus of CK1.2 are essential for the subcellular localisation of CK1.2 but not for its kinase activity, whereas the deletion of the N-terminus leads to a dramatic decrease in CK1.2 abundance. In conclusion, our data give new insights into the roles of *Leishmania* CK1.2 in the parasite.

## Results

### *Leishmania* CK1.2 localisation is ubiquitous

To investigate the localisation of CK1.2 in promastigotes, *Leishmania donovani* parasites expressing an episomal copy of CK1.2 tagged with V5-His_6_ at the C-terminus (CK1.2-V5) were used [13]. This cell line was validated in a previous study by showing that CK1.2-V5 is active, and functional as it compensates for a decrease of endogenous CK1.2 activity [13] indicating that ectopic CK1.2-V5 is properly folded. Immunofluorescence assays of parasites fixed with paraformaldehyde (PFA) were performed revealing intense punctate staining in the cytoplasm, the nucleus and the flagellum (Fig. 1A). The control parasites expressing the empty vector only showed weak background fluorescence (Fig. 1B). Indeed the sum of the fluorescence intensity in the cellular body of parasites expressing CK1.2-V5 (1 712 899 ± 85 178, n=256) was significantly higher than that of the control (893 556 ± 16 256, n=154) (Fig. 1C), suggesting that this punctate staining is specific to CK1.2-V5. This pattern is characteristic of the localisation of human CK1s [27]. As the cytoplasmic staining of CK1.2 could mask specific localisations, promastigotes were treated with 0,125% NP-40 to permeabilise membranes and facilitate the release of cytoplasmic proteins, prior to PFA fixation and staining. The treatment condition was selected after optimisation steps performed to minimise the impact of the detergent on the nucleus and kinetoplast (data not shown). We detected a signal (i) adjacent to the kinetoplast (Fig. 1D, CK1.2-V5) (ii) in the flagellar pocket region and along the flagellum and (iii) in the Hoechst-unstained region of the nucleus (Fig. 1D, CK1.2-V5). These signals were reproducibly observed in all the samples that were analysed. CK1.2-V5 specific localisation was confirmed using methanol treatment at different incubation times (as an example: 3 min, Figure S1A), ruling out the possibility that it could be an artefact of detergent treatment. To confirm these observations, co-localisation studies were performed with organelles-specific markers.

**FIGURE 1:**
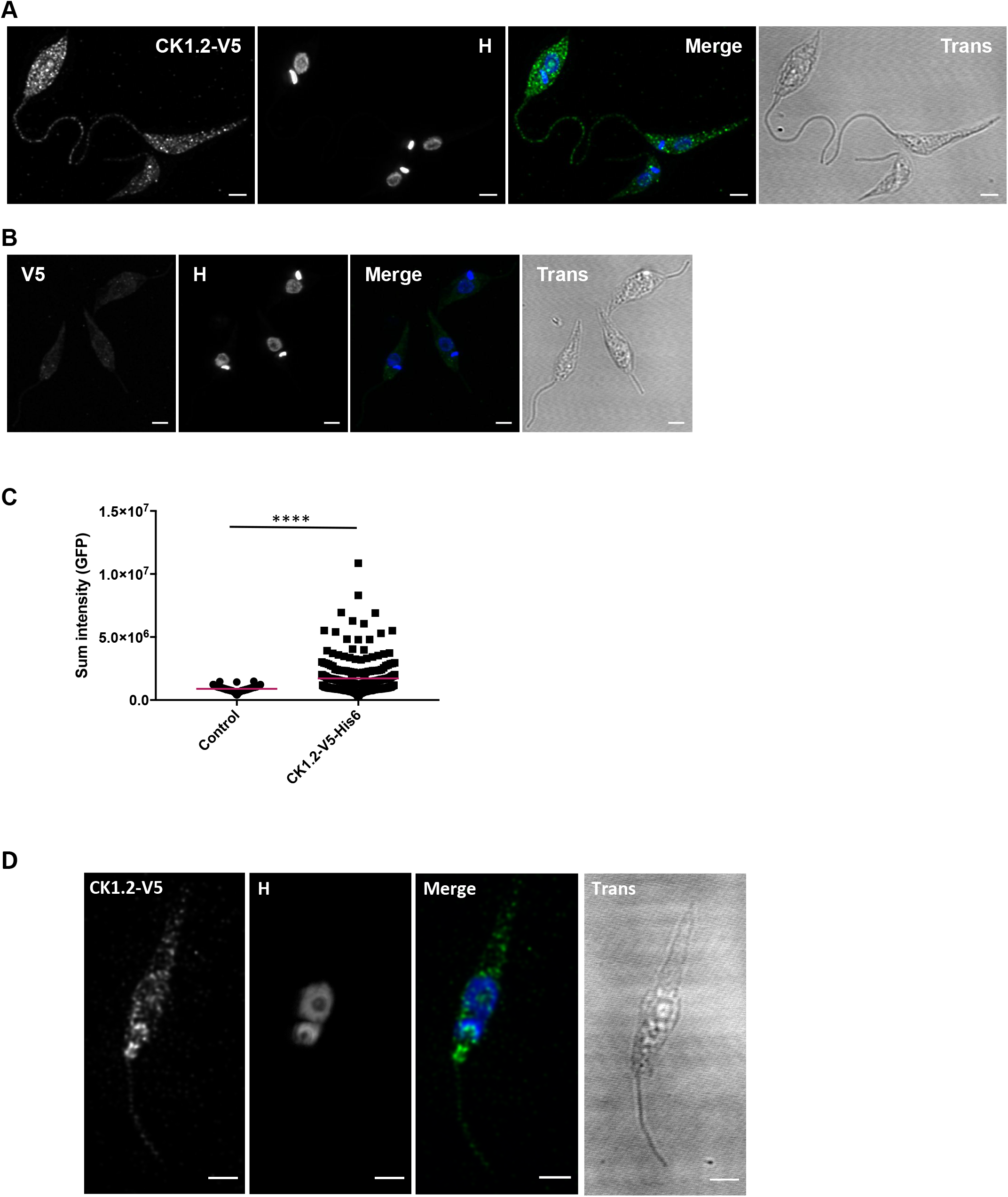
*Leishmania* CK1.2 localisation is ubiquitous. (A) IFA of *Ld*Bob pLEXSY-CK1.2-V5 and (B) *Ld*Bob pLEXSY (mock) promastigotes, fixed with PFA. The confocal images show the anti-V5 staining (CK1.2-V5 or V5, green), Hoechst 33342 staining (H, blue), a merge and the transmission image (Trans). Scale bar, 2 µm. The pictures are maximum intensity projection of the confocal stacks containing the parasites. (C) Analysis of different parameters extracted from ROI of the promastigotes parasite bodies. Scatter dot plots showing the sum of fluorescence intensity (for the V5 signal) of CK1.2-V5-expressing or mock control cell lines. The red line corresponds to the mean intensity. (D) IFA of *Ld*Bob pLEXSY-CK1.2-V5 promastigotes obtained after detergent treatment followed by PFA fixation. Similar to (A), except that the images are sum intensity projection of the confocal stacks containing the parasites.

### CK1.2 localises to the basal body and the flagellum

To investigate the localisation of CK1.2-V5 to the basal body [28], we used a specific marker, centrin-4 that localises to this organelle (Fig. 2A, CEN, white arrow) as well as to the bilobe structure (Fig. 2A, CEN, yellow arrow) [29] [30]. CK1.2-V5 co-localises with centrin-4 to the basal bodies (Fig. 2B panel a, white arrows) as measured by a mean Pearson coefficient (mPc of 0.741 ± 0.050 above 0.5, Fig. 2C), but not to the bilobe structure (Fig. 2B panel a, yellow arrow and Fig. 2C, mPc=0.47 ± 0.074). CK1.2 does not seem to have functions in the kinetoplast, as the kinase does not co-localise with the kinetoplast DNA (mPc=0.27 ± 0.164, Fig. 2C). Clearly, CK1.2 is not restricted to the basal bodies. Indeed, as judged by Figure 2B (panel b, white arrow) and confirmed by an mPc of 0,805 ± 0,06 (Fig. 2C), CK1.2-V5 co-localises with IFT172 at the transition fibers, suggesting that the protein kinase crosses the transition zone to enter into the flagellum. CK1.2-V5 localises to the axoneme similarly to IFT172 (Figure 2D, Merge and 3D-view), but not to the paraflagellar rod (PFR2, Fig. 2E, panel 3). These findings suggest that CK1.2 is perfectly located (i) to regulate basal body functions such as the coordination of kinetoplast/basal body segregation, and (ii) to cross the transition zone and access the flagellum.

**FIGURE 2:**
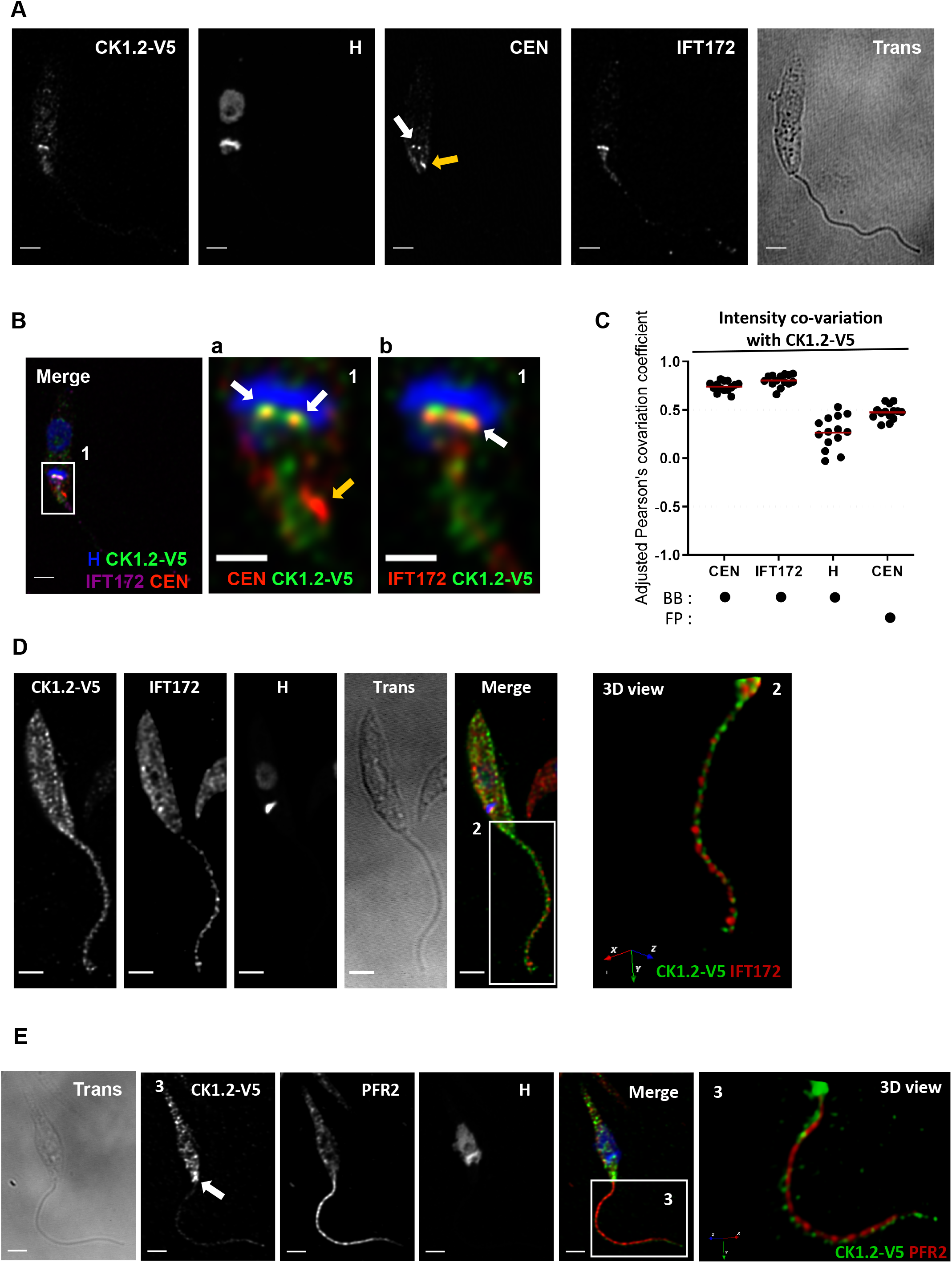
CK1.2 localises to the basal bodies and the flagellum. IFA of *Ld*Bob pLEXSY-CK1.2-V5 promastigotes obtained after detergent treatment followed by PFA fixation. (A) Single channel images of the CK1.2-V5, Hoechst 33342 (H), centrin 4 (CEN) or IFT172 signals and the transmission image (Trans). The white arrow highlights the basal bodies and the yellow arrow the bilobe region. (B) The merge panel shows CK1.2-V5 signal (green) merged with H (Hoechst 33342, blue), CEN (centrin 4, red) and IFT172 (cyan). The two right panels show a magnification of region 1 with H signal (blue) merged with (a) CK1.2-V5 (green) and CEN (red) and (b) IFT172 (red) and CK1.2-V5 (green) signals. Scale bar, 2 µm or 1 µm for magnified images. These pictures are single stacks extracted from deconvolved confocal stacks corrected for chromatic aberration. The white arrows highlight the basal bodies and the yellow arrow the bilobe region. (C) Dot plots showing Pearson’s covariation coefficients in the basal body (BB) or the flagellar pocket (FP) regions for different combination of signals. Pearson’s covariation coefficients were measured from n=14 different confocal stacks which were deconvolved and corrected for chromatic aberration with Huygens Professional software. The plot was generated with GraphPad Prism software and the mean values are represented with red bold segments. (D) IFA of *Ld*Bob pLEXSY-CK1.2-V5 promastigotes fixed by PFA and stained with the anti-V5 and anti-IFT172 antibodies. The left panels display the single channel images of the CK1.2-V5, IFT172, Hoechst 33342 (H) signals, the transmission image (Trans), and a merge of CK1.2-V5 (green), IFT172 (red) and H (blue) signals. The right panel shows a 3D-reconstruction (3D view) of the flagellum (image 2), with CK1.2-V5 signal (green) merged with IFT172 (red). (E) IFA of *Ld*Bob pLEXSY-CK1.2-V5 promastigotes obtained after detergent treatment followed by PFA fixation and staining with the anti-V5 and anti-PFR2 antibodies. The following images show CK1.2-V5, PFR2, Hoechst 33342 (H) signals, the transmission image (Trans), and finally the merge of CK1.2-V5 (green), PFR2 (red) and H (blue) signals. (image 3) 3D-reconstruction (3D view) of the flagellum (white square in (merge)), showing CK1.2-V5 signal (green) merged with PFR2 (red). Scale bars, 2 µm or 1 µm for 2D images. Pictures in (E) and (D, left panels) are single stacks extracted from deconvolved confocal stacks corrected for chromatic aberration. All confocal stacks containing the parasite were used for pictures (D, image 2 and E image 3).

### CK1.2 co-localises with Hsp90 and Hsp70 to the flagellar pocket

As shown Figure 2E (CK1.2-V5, white arrow), CK1.2-V5 is detected in the flagellar pocket (FP), which is important for processes such as exo/endocytosis or flagellum assembly [31]. The FP is also one of the sites of exosome excretion in *Trypanosoma brucei* [32]. Exosomes are vesicles of endosomal origin released by cells from multi-vesicular bodies into their extracellular environment and known to promote cell-to-cell communications [33]. CK1.2 was identified in *Leishmania* exosomes by proteome analyses, suggesting a role of this kinase in the host cell [24] [22]. Two other proteins are known to be exosomal protein cargos, Hsp90 and Hsp70. In human, these two chaperones are phosphorylated by human CK1 to control the balance between folding and degradation [34]. Likewise, *Leishmania* CK1.2 was recently shown to phosphorylate *Leishmania* Hsp90 [35] [34]. To investigate whether CK1.2 co-localises with Hsp90 and Hsp70 to the flagellar pocket, we performed immunofluorescence microscopy using detergent treatment to remove the cytoplasmic fraction of these chaperones.

#### CK1.2 co-localises with Hsp90 to the flagellar pocket neck

Most of Hsp90 was removed upon detergent treatment, indicating that the major fraction of Hsp90 is cytoplasmic (Fig. S2A). The small remaining pool of Hsp90 was associated with the flagellar pocket neck, where it co-localised with CK1.2-V5 (Fig. 3A, panel c) as confirmed by a mPc above 0.5 (Fig. 3B). The two proteins seem to form a horseshoe shaped structure as judged by the 3D view (Fig. 3C). These findings suggest that CK1.2 and Hsp90 may have specific functions linked to endo- or exocytosis, as the flagellar pocket neck is the site of endocytosis regulation [36]. Alternatively, they could be implicated in the regulation of the FAZ proteins, such as FAZ10 that has a similar localisation [37].

**FIGURE 3:**
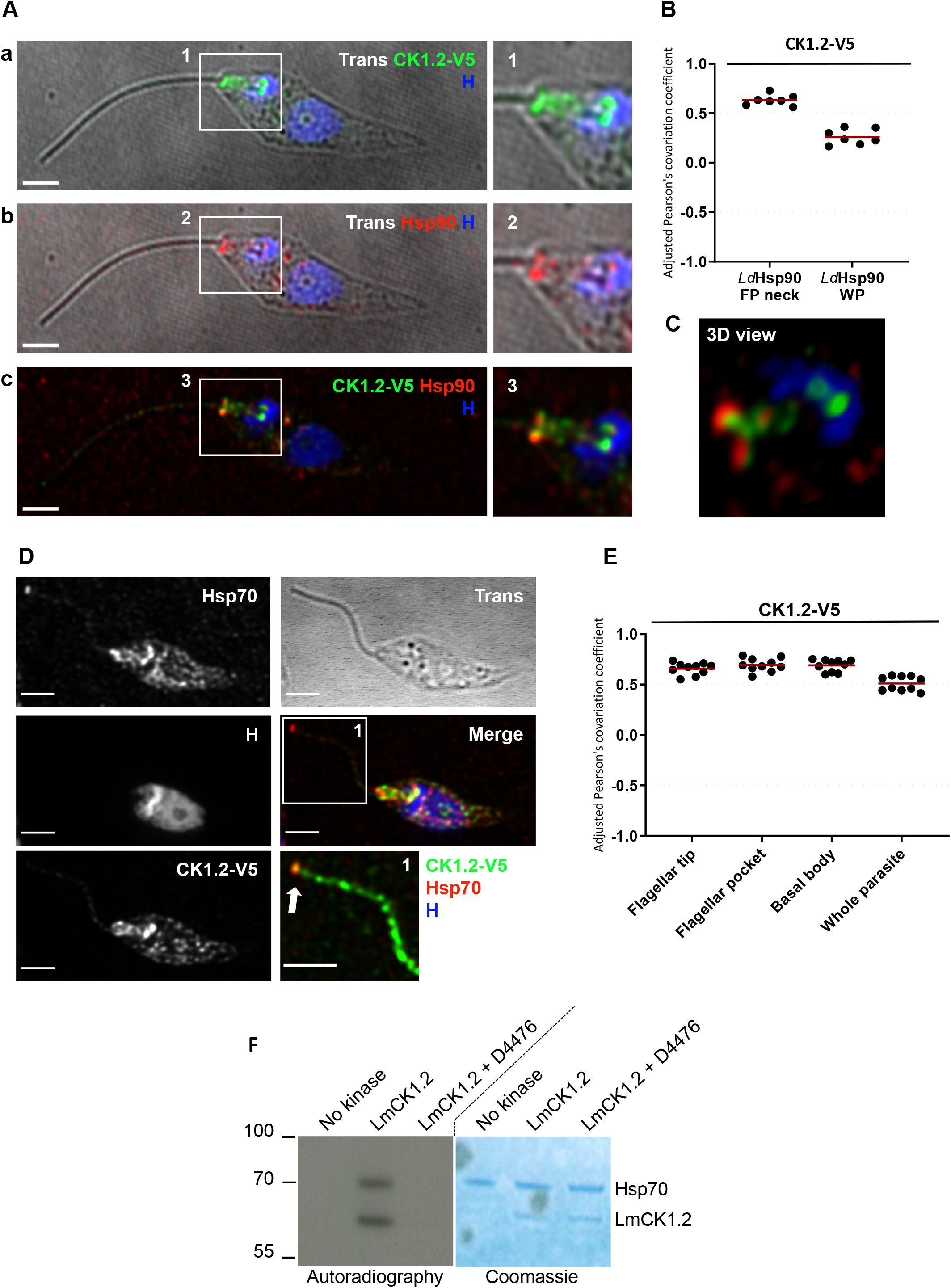
CK1.2 and Hsp90 co-localise to the flagellar pocket neck. Hsp70 co-localises with CK1.2 to the flagellum, to the flagellar tip and to the basal body. (A) IFA pictures of *Ld*Bob pLEXSY-CK1.2-V5 promastigotes obtained after detergent treatment followed by PFA fixation. (a and b) The left panel shows the transmission images merged with (a) Hoechst 33342 (H, blue) and CK1.2-V5 (green) signals, or (b) with H (blue) and Hsp90 (red) signals. The left panel in (c) shows a merged image of H (blue) with CK1.2-V5 (green) and Hsp90 (red) signals. The right panels show a magnification of the flagellar pocket and basal body region (white square) for their respective left panel. Scale bar, 2 µm. These pictures are single stacks extracted from deconvolved confocal stacks corrected for chromatic aberration. (B) Dot plots showing Pearson’s covariation coefficients for CK1.2-V5 and Hsp90 signals at the FPN or in the whole parasite region (WP). Pearson’s covariation coefficients were measured from n=7 different confocal stacks, which were deconvolved and corrected for chromatic aberration with Huygens Professional software. The plot was generated with GraphPad Prism software and the mean values are represented with red bold segments. (C) 3D-reconstruction of the anterior end of the parasite body from image (A) panel (c) (white rectangle region), showing CK1.2-V5 signal (green) merged with Hsp90 (red) and H (blue). All confocal stacks containing the parasite were used for this picture. (D) IFA pictures of *Ld*Bob pLEXSY-CK1.2-V5 promastigotes obtained after detergent treatment followed by PFA fixation. Single channel images show Hsp70, Hoechst 33342 (H, blue) and CK1.2-V5 (green) signals, and the transmission image (Trans). Region (1) shows a magnification of the flagellum (white square region) of the merged panel. The pictures are maximum intensity projection of the confocal stacks containing the parasites, after removal of the stacks in contact with the glass coverslip. Confocal stacks were deconvolved and corrected for chromatic aberration. Scale bar, 2 µm. The white arrow highlights Hsp70 and CK1.2 signal to the flagellar tip. (E) Dot plots showing Pearson’s covariation coefficients for CK1.2-V5 and Hsp70 signals in the flagellar tip, flagellar pocket, basal body regions or the whole parasite region. Pearson’s covariation coefficients were measured from n=10 different confocal stacks which were deconvolved and corrected for chromatic aberration with Huygens Professional software. The plot was generated with GraphPad Prism software and the mean values are represented with red bold segments. (F) *In vitro* kinase assay Hsp70. HSP70 was incubated with or without rCK1.2 and with rCK1.2 + D4476 (CK1 inhibitor) in presence of buffer C and γ-^32^P-ATP. Kinase assays were performed at 30°C for 30 min and reaction samples were separated by SDS-PAGE, gels were stained by Coomassie (right panel), and signals were revealed by autoradiography (left panel). The position of marker proteins is indicated on the left, the positions of CK1.2 and HSP70 are indicated on the right. Results are representative of two independent experiments.

#### CK1.2 co-localises with Hsp70 to the basal body, the flagellar pocket and flagellar tip and phosphorylates Hsp70

Cytoplasmic Hsp70 was removed upon detergent treatment (Fig. S2B and 3D). The remaining fraction of Hsp70 co-localises with CK1.2-V5 to the flagellum, the flagellar tip, the flagellar pocket, and the basal body, as confirmed by the mPc above 0.5 (Fig. 3D, merge and 3E). In fact, Hsp70 co-localises with CK1.2-V5 across the whole parasite (Fig. 3E, whole parasite), which was not observed for Hsp90 (Fig. 3B, WP). Hsp70 might thus be one of the main interactors of CK1.2. To investigate whether CK1.2 regulates Hsp70, kinase assays were performed using recombinant CK1.2-V5 and Hsp70 (Fig. 3F). The incorporation of ^32^P in Hsp70 in the presence of CK1.2-V5 indicates that Hsp70 is a substrate of CK1.2, similarly to its human orthologs. This result is further supported by the loss of phosphorylation following the addition of D4476, a specific inhibitor of CK1.2 [13]. We excluded the possibility that the phosphorylation was due to the ATPase activity of Hsp70, as no phosphorylation was detected in the absence of the kinase. These data suggest that Hsp70 may be regulated by CK1.2-mediated phosphorylation, explaining the co-localisation of both proteins.

### CK1.2 is localised in the granular zone of the nucleolus and redistributed to the mitotic spindle during mitosis

As shown Figure 1D, CK1.2-V5 was detected in a sub-nuclear location unstained by Hoechst, corresponding to the nucleolus. To ascertain this hypothesis, the localisation of CK1.2-V5 in detergent-treated promastigotes was compared to that of L1C6 antibody, which specifically recognises an unknown nucleolar protein [38]. The L1C6-targeted antigen was detected in the centre of the Hoechst-unstained area in the nucleus corresponding to the dense fibrillar zone of the nucleolus and thought to be involved in rDNA transcription (Fig. 4A, merged image-red staining; Fig. S3A) [39]. In contrast, CK1.2-V5 was detected at the periphery of the nucleolus, as dotted staining around L1C6 (Fig. 4A, merged image, green staining; Fig. S3A). This localisation corresponds to the granular component of the nucleolus, which contains mainly RNA and is thought to be involved in the last steps of rRNA processing and ribosome biogenesis. Thus, CK1.2 might be involved in rRNA processing rather than in rDNA transcription, which is a novel finding for CK1 family members. Furthermore, in dividing cells the staining of CK1.2 elongates from a wheel-shaped to a bar-shaped form that reaches both ends of the cell, similarly to the nucleolar region but unlike L1C6 staining (Fig. 4B, CK1.5-V5, H and L1C6; Fig. S3A). Instead, L1C6 antigen follows the classical segregation pattern described for nucleolar components (Fig. 4B, L1C6; Fig. S3A) [40]. As shown in Figure 4C panel a and Figure S3B, CK1.2-V5 also co-localises with the mitotic spindle in specific areas, observation confirmed by the mPc above 0.5 (Fig. 4D, mitotic spindle (reduced)). During anaphase, CK1.2-V5 is localised at each end of the elongated mitotic spindle (Fig. 4C panel d; Fig. S3B), similarly to the twinfilin-like protein [41]. These findings suggest that CK1.2 may be involved in cytokinesis or in the regulation of chromosome segregation [41]. Evidence from other eukaryotes support such a role for CK1 in mitosis and its recruitment to the spindle [42]. The mitotic spindle co-localises also with the nucleolus, suggesting that nucleolar proteins could be involved in chromosome segregation in the absence of visible centrosomes. Similar processes have been described in *Trypanosoma brucei* [43]. Thus, CK1.2 might be yet another nucleolar protein that relocates from the nucleolus to the mitotic spindle during mitosis.

**FIGURE 4:**
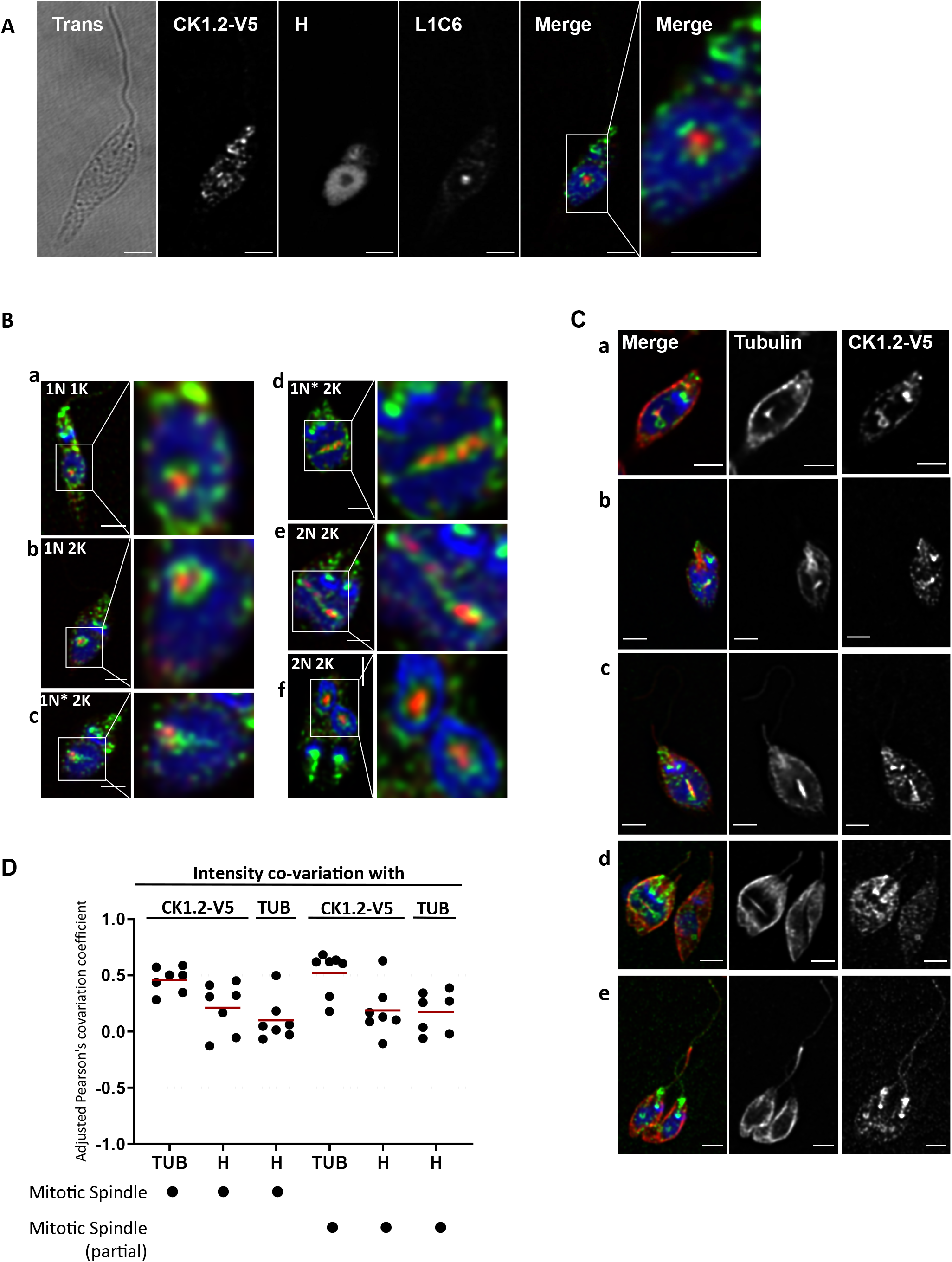
CK1.2 localises in the nucleolus and to the mitotic spindle. (A) IFA pictures of *Ld*Bob pLEXSY-CK1.2-V5 promastigotes obtained after detergent treatment followed by PFA fixation. The images show the single channel images of the transmission image (Trans), CK1.2-V5, Hoechst 33342 (H, blue) and L1C6 signals (red) and merged image. The right panel show a magnification of the nucleus region. Scale bar, 2 µm. These pictures are single stacks extracted from deconvolved confocal stacks corrected for chromatic aberration. (B) IFA pictures of *Ld*Bob pLEXSY-CK1.2-V5 promastigotes obtained after detergent treatment followed by PFA fixation and stained with anti-V5 (CK1.2-V5) and anti-L1C6 (nucleolus, L1C6) antibodies. Confocal images representing sequential events of mitosis revealed different localisation patterns of L1C6 nucleolar marker and CK1.2-V5. (a – f) The images show the merged image containing CK1.2-V5 (green), Hoechst 33342 (H) (blue) and L1C6 (red) signals and a magnification of the nuclear region. N=nucleus, K=kinetoplast. Scale bar, 2 µm or 1 µm for magnified images. These pictures are single stacks extracted from deconvolved confocal stacks corrected for chromatic aberration. See also Figure S3A. (C) IFA pictures of *Ld*Bob pLEXSY-CK1.2-V5 promastigotes obtained after detergent treatment followed by PFA fixation and stained with anti-V5 and anti-α-tubulin antibodies. Sequential images of various stages of cell division (a – e) showing the single channel images for α-tubulin signals and the merged images showing CK1.2-V5 (green), H (blue) and α-tubulin (red) signals. Scale bar, 2 µm. These pictures are single stacks extracted from deconvolved confocal stacks corrected for chromatic aberration. See also Figure S3B. (D) Dot plots showing Pearson’s covariation coefficients for different combination of signals in the entire mitotic spindle region and in a reduced mitotic spindle region containing also CK1.2-V5 signal. For both regions, CK1.2-V5 signal was compared with α-tubulin (TUB) or Hoechst 33342 (H). The signal of TUB was also compared with H. Pearson’s covariation coefficients were measured from n=7 different confocal stacks which were deconvolved and corrected for chromatic aberration with Huygens Professional software. The plot was generated with GraphPad Prism software and the mean values are represented with red bold segments.

### CK1.2 has a similar localisation in axenic amastigotes than in promastigotes

In PFA-fixed axenic amastigotes, the localisation of CK1.2-V5 is similar to that observed in promastigotes with intense fluorescent dots in the cytoplasm and at the flagellar tip (Fig. 5A, white arrows). In detergent treated or untreated axenic amastigotes, CK1.2-V5 localises to similar structures as observed in promastigotes (Fig. 5B). In contrast to untreated axenic amastigotes, in detergent-treated axenic amastigotes CK1.2-V5 seems to be excluded from the flagellar tip and restricted to the flagellar pocket neck, where it forms a horseshoe-shaped structure as judged by Figure 5C (panel b, green staining). Indeed, in PFA, CK1.2 was detected at the flagellar tip of 82% of CK1.2 positive cells, whereas in detergent, it was detected in only 8% of CK1.2 positive cells. This result suggests that CK1.2 is not associated with the cytoskeleton at the flagellar tip in contrast to what has been observed in promastigotes.

**FIGURE 5:**
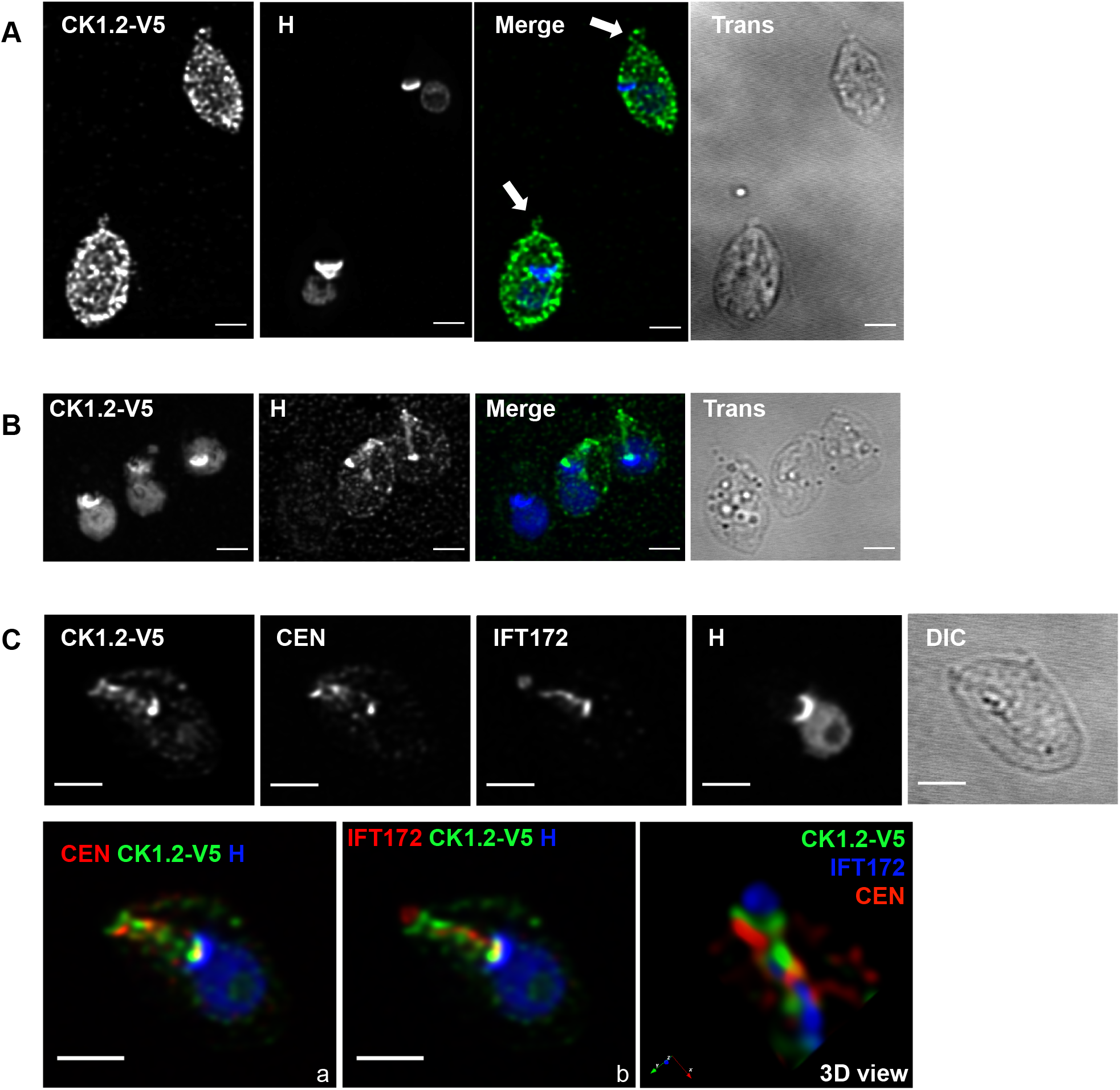
CK1.2 localisation in amastigotes. IFA of *Ld*Bob pLEXSY-CK1.2-V5 axenic amastigotes, fixed with PFA (A) or obtained after detergent treatment (B). The single channel images show CK1.2-V5 (green) and Hoechst 33342 (H, blue) signals, and the transmission image (Trans). Scale bar, 2 µm. The pictures are maximum intensity projection of the confocal stacks containing the parasites. Confocal stacks were deconvolved and corrected for chromatic aberration. (C) IFA of *Ld*Bob pLEXSY-CK1.2-V5 axenic amastigotes obtained after detergent treatment followed by PFA fixation. The single channel and the merge images show CK1.2-V5, Hoechst 33342 (H, blue), centrin 4 (CEN, red) or IFT172 (cyan) signals and the transmission image (Trans). Scale bar, 2 µm. These pictures are single stacks extracted from deconvolved confocal stacks corrected for chromatic aberration. Panel a shows a merged of CK1.2- V5, Hoechst 33342 (H, blue) and centrin 4 (CEN, red); Panel b shows CK1.2-V5, Hoechst 33342 (H, blue), and IFT172 (cyan) signals; 3D-reconstruction of the flagellar pocket region and its neck from image (C), which has been rotated. All confocal z-stacks containing the parasite were used for the 3D view.

CK1.2 displays multiple localisation patterns, which are likely to be associated with pleiotropic functions. How *Leishmania* CK1.2 is targeted to these different localisations remains to be investigated, especially considering that there are no specific motifs, apart from a non-functional nuclear localisation signal [1]. Furthermore, *Leishmania* CK1.2 is constitutively active, contrary to human CK1δ, ε and to a lesser extent CK1α, thus probably requires tighter regulation mechanisms to avoid inappropriate phosphorylation of its substrates [13] [1].

### The low complexity regions are essential for *Leishmania* CK1.2 localisation

In higher eukaryotes, the activity and localisation of CK1 are mainly regulated through its N- and C-terminal domains [1, 44]. Among *Leishmania* CK1 paralogs, CK1.2 and CK1.1 are highly similar but differ in their N- and C-terminus, which might explain the differences in regulation and localisation [13, 45]. In contrast to CK1.1, CK1.2 is released into the host cell via exosomes and is essential for parasite survival [13, 22, 24]. N- and C-terminal truncations of CK1.2 were thus generated, based on the alignment with CK1.1 to determine the importance of these domains for the localisation and regulation of CK1.2 [45]. As shown in Figure 6A, the three truncated CK1.2 proteins were (i) lacking the last ten amino acids (aa) at the C-terminus (CK1.2ΔC10), (ii) lacking the last 43 aa at the C-terminus (CK1.2ΔC43), or (iii) lacking the first seven aa at the N-terminus (CK1.2ΔN7). To test whether these mutants were still active kinases [44], recombinant mutants proteins were expressed and used to perform a kinase assay with MBP, a canonical substrate for CK1.2 [13]. CK1.2, CK1.2ΔC10, CK1.2ΔC43 and CK1.2ΔN7 were equally active, as demonstrated by the incorporation of ^32^P into MBP (Fig. 6B, top panel). This finding indicates that the N- or the C-terminal domains are not essential for the activity of CK1.2. Next, promastigotes were transfected with a pLEXSY plasmid empty (control) or containing CK1.2, CK1.2ΔC10, CK1.2ΔC43, CK1.2ΔN7 genes. The expression of the three truncated proteins was analysed by Western blot analysis (Fig. 6C). CK1.2ΔC10-V5 and CK1.2ΔC43-V5 levels were similar to that of CK1.2-V5, whereas the level of CK1.2ΔN7 was lower (Fig. 6C). At least two hypotheses could explain the low abundance of CK1.2ΔN7, either the deletion of the N-terminus leads to structural instability or to degradation. We excluded the first possibility, since CK1.2ΔN7 was easily produced as an active recombinant kinase in bacteria (Fig. 6B, bottom panel). To test the second hypothesis, transgenic parasites expressing CK1.2ΔN7 were treated with Mg132, a proteasome inhibitor. The level of CK1.2ΔN7 as well as that of CK1.2 and the other mutants were similar in presence or absence of Mg132 (Fig. S4A panel a). The treatment of mutants was sufficient to block proteasomal degradation, as the level of ubiquitinated proteins was increased in presence of Mg132 (Figure S4A panel b). We next investigated whether the CK1.2ΔN7 may be degraded in the lysosomes. To this end, transgenic parasites were treated with ammonium chloride (NH_4_Cl), which increases the pH in the lysosome rendering hydrolase inactive [46]. The level of CK1.2ΔN7 remains low (Figure S4B panel a) despite the inhibition of lysosomal proteases as judged by the alkalinisation of the lysosome by NH_4_Cl and by the decrease in LysoTracker fluorescence intensity, which stains acidic compartments (Figure S4B panel b). The low level of CK1.2ΔN7 protein is thus not the consequence of proteasomal, lysosomal degradation or autophagy, which is ultimately a lysosome-mediated degradation [47]. Cathepsin B- or calpain-like cysteine peptidase-mediated degradation were excluded, as Mg132 also inhibits these proteases [48].

**FIGURE 6:**
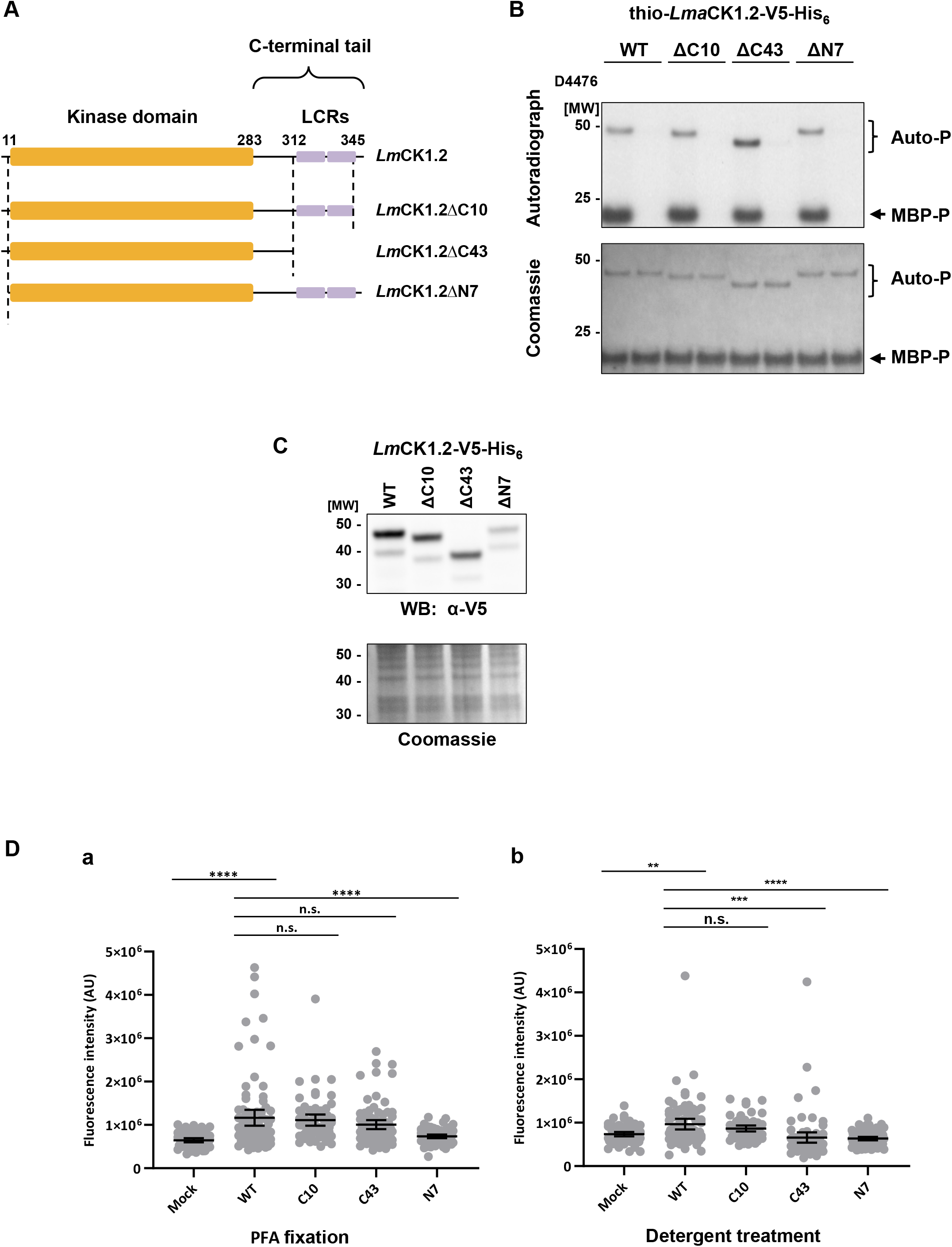
The C-terminal domain of CK1.2 is essential for its localisation to specific organelles, but not for its activity. (A) Cartoon representing the domain structure of *Lm*CK1.2, CK1.2ΔC10, CK1.2ΔC43 and CK1.2ΔN7 (GenBank: CBZ38008.1). The protein contains a kinase domain (yellow) and a C-terminal tail with two low complexity regions (LCR) (purple). (B) *In vitro* kinase assay using recombinant thio-CK1.2- V5 (WT, 55.9 kDa) and the truncated kinase mutants thio-CK1.2-ΔC10-V5 (ΔC10, 54.9 kDa), thio-CK1.2-ΔC43-V5 (ΔC43, 52.1 kDa) and thio-CK1.2-ΔN7-V5 (ΔN7, 55.2 kDa). Results are representative of three independent experiments. Purified proteins were incubated with MBP as substrate, with or without D4476. Kinase assays were performed for 30 min at pH 7.5 and 30°C and reaction samples were separated by SDS-PAGE, gels were stained by Coomassie (bottom), and signals were revealed by autoradiography (top). The brackets indicate auto-phosphorylation (Auto-P) and the arrows substrate phosphorylation (MBP-P) signals. MW= Molecular Weight. (C) Western blot analysis. Proteins were extracted from *Ld*Bob pLEXSY-CK1.2-V5 (WT, 42.8 kDa) or expressing truncated kinase mutants *Ld*Bob pLEXSY-CK1.2-ΔC10-V5 (ΔC10, 41.8 kDa), *Ld*Bob pLEXSY-CK1.2-ΔC43-V5 (ΔC43, 39.0 kDa) and *Ld*Bob pLEXSY-CK1.2-ΔN7-V5 (ΔN7, 42.1 kDa) in logarithmic phase promastigotes. Twenty micrograms were analysed by Western blotting (WB) using the anti-V5 antibody (α-V5) (top panel). The Coomassie-stained membrane of the blot is included as a loading control (bottom panel). MW= Molecular Weight. The blot is representative of three independent experiments. (D) Measurement of the sum of fluorescence intensity extracted from ROI of the promastigote parasite bodies in the mock, wild type and three domain-deletion mutants (same cell lines as in (C)). Scatter dot plots showing the sum fluorescence intensity (V5 signal) in the different cell lines in PFA-fixed (a) or detergent-treated (b) parasites. Data originates from n=54 (Mock, PFA), n=88 (WT, PFA), n=63 (ΔC10, PFA), n=80 (ΔC43, PFA), n=68 (ΔN7, PFA), n=62 (Mock, det. treated), n=77 (WT, det. treated), n=55 (ΔC10, det. treated), n=78 (ΔC43, det. treated) and n=74 (ΔN7, det. treated). The mean values and the 95% confidence intervals are indicated with bold segments. Statistically significant differences are indicated with two (p < 0.01), three (p < 0.001) or four asterisks (p < 0.0001). ns. = non-significant.

To assess the importance of the N- and C-terminal domains for CK1.2 localisation, immunofluorescence studies were performed either on PFA-fixed cells or on detergent-treated PFA-fixed cells as previously described. To take into consideration the heterogeneity of CK1.2-V5 staining (Fig. 1C), the sum fluorescence of each parasite was measured and that of the WT was compared to that of the three mutant parasites. No statistically significant differences could be measured between CK1.2, and CK1.2ΔC10 or CK1.2ΔC43, in PFA-fixed cells (Fig. 6D panel a). In contrast, a statistically significant difference was measured between CK1.2 and CK1.2ΔN7. This result is consistent with the data obtained from the Western blot analyses (Fig. 6C) and suggest that the C-terminal deletions do not decrease the level of the kinase in the parasite. Next, the same experiment was performed using detergent-treated parasites to evaluate the ability of the mutant proteins to associate with the cytoskeleton or to localise to organelles (Fig. 6D panel b). There was no significant difference in fluorescence intensity between CK1.2 and CK1.2ΔC10, suggesting that the last 10 amino acids are not required for the specific localisation of CK1.2. Conversely, a significant difference in fluorescence intensity was measured between CK1.2 and CK1.2ΔN7, which was expected, and between CK1.2 and CK1.2ΔC43, indicating that the level of the two mutant proteins detected in the cells after detergent treatment was lower than that of the WT. The total level of CK1.2ΔC43 did not change (Fig. 6C and 6D panel a), only the fraction associated with specific organelles or the cytoskeleton was reduced, suggesting that the decrease in intensity is the consequence of a lack of proper localisation of this mutant protein. In summary, ours results suggest that the last 10 aa at the C-terminus are not implicated in the subcellular localisation of CK1.2, in contrast to the domain between aa 310 and 343, which corresponds to the low complexity regions absent in CK1.1. Deleting this domain prevents CK1.2 from associating with organelles and the cytoskeleton, thus it remains in the cytoplasm.

## Discussion

Although, CK1.2 is essential for promastigotes, axenic and intra-macrophagic amastigotes, little is known about the essential functions it performs in the parasite and in the host cell [13]. The data presented here show that CK1.2 displays a pleiotropic localisation, consistent with its involvement in multiple processes. This finding is similar to the data obtained with its orthologs [1]. As shown in higher eukaryotes, localisation of CK1 is linked to its functions and regulates its specificity towards its substrates. Thus, the localisation of *Leishmania* CK1.2 provides insights into its functions [26], especially since the localisation or functions of CK1.2 orthologs in other parasites is largely unknown. In *Plasmodium falciparum*, the localisation of PfCK1 depends on the life cycle and is mainly at the surface of the red blood cells in early stages of infection and restricted to the parasite in mature trophozoites and merozoites [12]. In *Toxoplasma gondii*, the localisation is cytoplasmic [49]. Because *Leishmania* CK1.2 has 73% identity to TbCK1.2, 85% to TcCK1.2, 69% to TgCK1a and 62 % to PfCK1 [13], our data are transferable to other parasites.

### Localisation of CK1.2 and its possible functions

Based on the localisation of CK1.2 and compared to the localisation and functions of its orthologs, several hypotheses could be made on its potential functions. *Leishmania* CK1.2 was detected in punctate structures in the cytoplasm of promastigotes as well as amastigotes. This localisation is characteristic of the CK1 family [27]. Although, the origins of these structures are unknown in *Leishmania*, human CK1δ and HRR25, its *Saccharomyces cerevisiae* ortholog are known to localise to P-bodies, which protect the kinase from degradation, especially during stress [11, 50]. P-bodies also store repressed mRNAs that mainly encode for regulatory processes [50]. Trypanosomatids contain P-bodies as well as other granules such as stress and heat shock granules [51] [52] [53], suggesting that such a localisation is conceivable for *Leishmania* CK1.2. Indeed, *Trypanosoma brucei* CK1.2 was recently shown to regulate ZC3H11, a protein involved in the stabilisation of stress response mRNAs [54]. This protein is mainly localised in the cytoplasm [55], which is consistent with the localisation of CK1.2 and might suggest a role in mRNA stabilisation for the *Leishmania* kinase.

CK1.2 is localised to the basal body, similarly to human CK1ε, which was shown to be involved in primary cilia disassembly [56], and to the axoneme, consistent with proteomic data that identified CK1.2 among the flagellar proteins in *T. brucei* and *Leishmania mexicana* [57] [58]. With *Chlamydomonas reinhardtii* and *Trypanosoma brucei*, *Leishmania* is the only eukaryote showing a flagellar localisation of CK1 [59]. These findings suggest that CK1.2 could be involved in motility [59].

The localisation of CK1.2 at the flagellar pocket suggests that the kinase might be exported by and/or regulate endocytosis. There are evidences supporting the two hypotheses: (i) CK1.2 is exported by exosomes, probably through the FP [22] [32]; and (ii) Hrr25, as well as human CK1δ/ε promotes initiation of clathrin-mediated endocytosis through its recruitment to endocytic sites [10]. We showed for the first time that Hsp90 is located at the flagellar pocket and more specifically to the neck, where it co-localises with CK1.2. Because CK1.2 phosphorylates Hsp90, both proteins may be involved in functions associated with the FPN such as facilitating the entry of macromolecules or regulating endocytosis [35, 36, 60]. However, given the localisation of these two proteins, an involvement in the regulation of FAZ proteins cannot be excluded and will be further investigated. In human cells, the phosphorylation of Hsp90 by human CK1 was shown to regulate the balance between protein folding and degradation [34]. CK1.2 also shares a cell-wide distribution with Hsp70, suggesting that both proteins might interact. Here, we demonstrated that Hsp70 is a substrate of CK1.2, which is consistent with the phosphorylation of human Hsp70 by human CK1 [34]. The results from the kinase assay and the co-localisation studies suggest that Hsp70 might be involved in the regulation of CK1.2 localisation. The roles of these complexes are unknown but should be explored further as Hsp70, Hsp90 and CK1.2 are exported via exosomes [22].

CK1.2 was detected in the nucleus and more specifically in the nucleolus where it seems to have a dual function. It might be involved in the regulation of the last steps of ribosomal processing rather than in rDNA transcription. This is different from previous data related to yeast and human CK1s, implicating them in the maturation of pre-40S ribosomes in the cytoplasm [61] [62]. Nevertheless, human CK1α and δ have been identified in the proteome of the nucleolus, suggesting that, similarly to *Leishmania* CK1.2, they might play a role in this organelle [63]. The second function of nucleolar CK1.2 might be linked to chromosome segregation. Indeed, we showed that nucleolar CK1.2 co-localises with tubulin from the assembly of the mitotic spindle to its elongation. This is consistent with the nucleolus as a site of mitotic spindle elongation and thus of chromosome segregation [41] [64]. Our data suggest that the nucleolar pool of CK1.2 might be redistributed onto the mitotic spindle during mitosis, and by analogy to human CK1α might be involved in spindle positioning [65]. The redistribution of nucleolar proteins has been described for other kinetoplastid proteins such as TbNOP86, a protein potentially involved in chromosome segregation in *T. brucei* [43] and LdTWF, an actin-binding protein that controls mitotic spindle elongation in *Leishmania* [41]. The knockdown of TbCK1.2 in bloodstream form parasites generates multinucleated cells [66]. These findings are consistent with a role of *Leishmania* CK1.2 in kinetoplast and chromosome segregation.

### Regulation of CK1.2 localisation

The subcellular localisation of mammalian CK1 depends on interacting partners such as FAM83 proteins [1, 27], suggesting that the N- and C-terminal domains, which are involved in protein-protein interactions, are crucial for the regulation of CK1 localisation. However, nothing was known about the motifs important to drive these interactions. Here, we showed that the C-terminal domain of CK1.2, between the amino acids 310 and 343, contains two Low Complexity Regions (LCRs), which are essential for its localisation. Indeed, LCRs were shown to be more abundant in highly connected proteins, such as signalling kinases [67], and may contribute to the binding of interacting partners. Removing these domains reduced the ability of CK1.2 to localise to specific organelles and subcellular structures. These LCRs are absent from the C-terminus of *Leishmania* CK1.1; instead one LCR is found at the N-terminus [68]. Based on the hypothesis that the LCRs drive the specificity of protein interactions, the localisation of CK1.1 should be different from that of CK1.2. Indeed, CK1.1 is a low abundant protein that was not detected in the flagellum, the basal bodies, the nucleolus, or the mitotic spindle [45]. Moreover, CK1.1, unlike CK1.2, was not identified as an exosomal cargo. The presence of LCRs in the C-terminus, which is the main difference between the two proteins, could thus be essential for the export of CK1.2 and its potential functions in the host cell [22, 24] [69]. Consequently, the identification of CK1.2 binding partners will be critical to understand how CK1.2 localisation is regulated. These LCRs are also found in the C-terminal domain of CK1α, CK1δ and CK1ε, as well as in the C- and N-terminus of CK1γ1, CK1γ2, CK1γ3, suggesting that they might share similar characteristics to that of *Leishmania* CK1.2 and thus be crucial for their localisations (Fig. S5, http://smart.embl-heidelberg.de/smart/set_mode.cgi?NORMAL=1, [70]). Recently, Fulcher et *al.* identified the FAM83A-H protein family, which act as subcellular anchors for CK1 isoforms through the conserved N-terminal domain of unknown function 1669 (DUF1669). FAM83 proteins could be part of an important mechanism for targeting CK1 activity to specific subcellular locations and substrates [27]. There are no orthologs in *Leishmania* as judged by protein alignment but similar mechanisms seems nevertheless to exist.

Likewise, the N-terminus is important for CK1.2, although its role remains elusive. Even though the removal of the N-terminus renders the protein undetectable in the cell, a degradation of CK1.2 by the proteasome, the lysosome or by autophagy was excluded. We hypothesise that the protein is directly excreted into the extracellular space [48]. This hypothesis is supported by the fact that *Leishmania* CK1.2, CK1.4 and *Plasmodium* CK1 were shown to be shedded into the extracellular medium [23, 71] [12]. Interestingly, the seven N-terminal amino acids are naturally absent in CK1.1, consistent with this protein being hardly detectable by Western blot or microscopy [45]. Remarkably, in contrast to several CK1 orthologs in eukaryotes, the N- and C-termini are not required for kinase activity. Indeed CK1 was shown to be inhibited by auto-phosphorylation, thus truncation of its C-terminus increases kinase activity [1]. We do not observe this phenomenon with L-CK1.2, suggesting that the kinase might not be auto-inhibited by auto-phosphorylation, similarly to mammalian CK1s.

In conclusion, we provide the first insight into the localisation and the potential associated functions of CK1.2. The identification of the interacting proteins that drive the localisation of CK1.2 to these different organelles will be instrumental for the precise characterisations of its functions in *Leishmania* as well as in other parasites. Moreover, our present and previous data demonstrate the similarity of localisation, structure, activity, regulation between *Leishmania* CK1.2 and human CK1s, highlighting that *Leishmania* CK1.2 is an excellent model to study mammalian CK1s (these data, [72] [13] [73]). Indeed, we uncovered novel localisations of CK1 family members including the nucleolus and our data provide the first analysis of CK1 regulatory domains in parasites and the first demonstration of the importance of LCRs for CK1 localisation in eukaryotes. Finally, to date CK1.2 is the only *Leishmania* signalling kinase shown to be exported into the host cell via exosomes and to have the ability to regulate multiple host cell processes, suggesting that it could be a key player for host-pathogen interactions.

## Materials and Methods

### Leishmania cell lines

All the parasite cell lines used in this study were derived from *L. donovani* axenic 1S2D (MHOM/SD/62/1S-CL2D) clone LdBob, obtained from Steve Beverley, Washington University School of Medicine, St. Louis, MO. Promastigotes were cultured and differentiated into axenic amastigotes as described previously [45]. Parasites cell lines were grown in media with 30 µg/mL hygromycin B (ThermoFisher Scientific Cat# 10687010) to maintain the pLEXSY-CK1.2-V5 or the empty pLEXSY plasmids. The transgenic *L. donovani* cell lines containing either the pLEXSY or pLEXSY-CK1.2-V5-HIS_6_ (pLEXSY-CK1.2-V5) vectors, corresponding to the mock or expressing *Leishmania major* CK1.2 tagged with V5 and HIS_6_, respectively, were described previously [13].

### Plasmids

For the generation of CK1.2ΔC10-V5-His_6_-, CK1.2ΔC43-V5-His_6_- and CK1.2ΔN7-V5-His_6_-expressing cell lines, we first amplified the V5-His_6_ fragment from pBAD-thio-topo-CK1.2 [13] using the following primers: 5’-gatggcattctagaatcgatgatatccccgggggtaagcctatcc-3’ and 5’- gcatggatcgcggccgctcaatggtg-3’. Then we digested the PCR fragment with XbaI and NotI and cloned it into the pLEXSY-Hyg plasmid (Jena bioscience) digested with the same enzymes to obtain pLEXSY- V5-His_6_ plasmid. Next, we amplified CK1.2ΔC10, CK1.2ΔC43 and CK1.2ΔN7 from pLEXSY-CK1.2 [13] with the following primers for CK1.2ΔC10: 5’-gatggcatcggatccatgaacgttgagctgcgtgt-3’ and 5’- gcatggatctctagagtttgcgctgttcggagc-3’; for CK1.2ΔC43: 5’-gatggcatcggatccatgaacgttgagctgcgtgt-3’ and 5’-gcatggatctctagagctttgctgttcctgcag-3’; for CK1.2ΔN7: 5’-gatggcatcggatccatgggtaatcgctatcgtattgg-3’, 5’-gcatggatctctagattgttgttccggtgcgccg-3’. We digested the PCR fragments with BglII and XbaI and cloned them into the pLEXSY-V5-His_6_ digested with the same enzymes to obtain respectively pLEXSY-CK1.2ΔC10-V5-His_6,_ pLEXSY-CK1.2ΔC43-V5-His_6_ and pLEXSY-CK1.2ΔN7-V5-His_6_. Finally, these vectors were transfected in LdBob. We generated *E.coli* strains containing pBAD-thio-topo-LmaCK1.2ΔC10-V5-His_6_, pBAD-thio-topo-LmaCK1.2ΔC43-V5-His_6_ or pBAD-thio-topo-LmaCK1.2ΔN7-V5-His6 by amplifying the whole pBAD-thio-topo-LmaCK1.2-V5-His_6_ except the last 30 bp, the last 129 bp or the first 18bp, respectively, using the following primers for CK1.2ΔC10: His_6_ *5’-AAGGGCGAGCTTGAAGGTAAG-3’* and *5’-GTTTGCGCTGTTCGGAGCG-3’*; for CK1.2ΔC43: *5’-AAGGGCGAGCTTGAAGGTAAG-3’ and 5’-GAAGCTTTGCTGTTCCTGC-3’*; and for CK1.2ΔN7: *5’-GGTAATCGCTATCGTATTGGTC-3’ and 5’-CATAAGGGCGAGCTTGTCATC-3’*. The linear PCR products were circularised by ligation with T4 DNA ligase (Promega Cat#M180A) (O/N, 4°C). Finally the plasmids pBADthio-*Lma*CK1.2ΔC10-V5-His_6_, pBADthio-*Lma*CK1.2ΔC43- V5-His_6_ and pBADthio-*Lma*CK1.2ΔN7-V5-His_6_ were sequenced and transformed in *Escherichia coli* Rosetta (DE3) pLysS Competent Cells (Merck Cat# 70956) for bacterial expression.

### Immunofluorescence

Logarithmic phase promastigotes or axenic amastigotes (48h after shift at 37°C and pH5.5) were resuspended at 2×10^6^ parasites per mL in Dulbecco’s Phosphate Buffer Saline (DPBS) (Gibco) and 500 µL were added to poly-L-lysine-coated coverslips placed in a 24-well plate. Plates were centrifuged 10 min at 1200 g at room temperature to settle parasites onto the coverslips. For fixation alone, cells were washed three times with DPBS and fixed in 4% paraformaldehyde (PFA) in DPBS for 15 min at room temperature. For cytoskeleton preparation, the protocol was adapted from [74]. Briefly, cells were washed three times with DPBS, treated with 0.125% Nonidet 40 (Fluka BioChemika Cat# 74385) in PIPES buffer (100 mM piperazine-N,N-bis(2-ethanesulfonic acid) (PIPES) pH6.8, 1 mM MgCl_2_) for 2 minutes at room temperature and washed twice for 5 minutes in PIPES buffer. Cells were fixed in 4% PFA in DPBS for 15 min at room temperature. After PFA fixation, cells were washed three times in DPBS, neutralised 10 min with NH_4_Cl (50 mM in DPBS), and washed again three times in DPBS. For the immuno-labelling of PFA-fixed cells or cytoskeleton preparations, the samples were blocked with 10% filtered heat-inactivated fetal calf serum (FCS) containing 0.5 mg.mL^-1^ saponin in DPBS for 30 min at room temperature and then washed for 5 min in DPBS. The cells were then incubated with primary antibodies diluted in DPBS with 0.5% Bovine Serum Albumin (BSA) and 0.5 mg.mL^-1^ saponin for 1h at room temperature. Three washes of 10 min were performed and the secondary antibody diluted in DPBS with 0.5% BSA and 0.5 mg.mL^-1^ saponin was added. After one hour incubation at room temperature in the dark, cells were washed twice for 10 min in DPBS with 0.5% BSA and 0.5 mg.mL^-1^ saponin, and then twice in DPBS. Parasites were incubated with 5 µg.mL^-1^ Hoechst 33342 in DPBS for 8 min in the dark, washed twice with DPBS, one time with distilled water, air-dried, then mounted with slides using SlowFade Gold Antifade Mountant (ThermoFisher Scientific Cat# S36937). For methanol fixation, logarithmic phase promastigotes were washed twice in DPBS and resuspended at 2×10^7^ parasites per mL. 10^6^ parasites were spread onto poly-L-lysine coated slides, and allowed to settle for 30 min in a humid chamber. Parasites were then fixed in methanol at −20°C for 3 minutes and rehydrated for 10 min in DPBS at room temperature. For immuno-labelling of methanol-fixed parasites, samples were blocked with 10% filtered heat-inactivated FCS in DPBS for 15 min at room temperature and washed for 5 min in DPBS. Then the cells were treated similarly as those fixed by PFA. The antibodies used were: mouse IgG2a anti-V5 tag monoclonal antibody (Thermo Fisher Scientific Cat# R960-25, RRID:AB_2556564) diluted at 1/200 (in PFA and methanol fixed parasites) or at 1/300 (in cytoskeleton preparations); rabbit anti-V5 tag polyclonal antibody (Abcam Cat# ab9116, RRID:AB_307024) diluted at 1/400; rabbit anti-LdCentrin polyclonal antibody (kind gift from Hira L. Nakhasi) diluted at 1/2000 [75]; mouse IgG1 anti-IFT172 monoclonal antibody diluted at 1/200 [76]; mouse anti-PFR2 L8C4 clone antibody diluted at 1/10 [77]; mouse L1C6 anti-TbNucleolus monoclonal antibody diluted at 1/100 (kind gift from Keith Gull [78]); mouse IgG1 anti-α-tubulin monoclonal DM1A antibody (Sigma-Aldrich Cat# T9026, RRID:AB_477593) diluted at 1/400; chicken anti-Hsp70 and chicken anti-Hsp90 antibodies diluted at 1/200 [79]. IgG subclass-specific secondary antibodies coupled to different fluorochromes were used for double labelling: anti-mouse IgG (H+L) coupled to AlexaFluor488 (1/200 (PFA- or methanol-fixed parasites)) or 1/300 (cytoskeleton preparations), Thermo Fisher Scientific Cat# A-21202, RRID:AB_141607)); anti-mouse IgG2a coupled to Cy3 (1/600; Jackson ImmunoResearch Labs Cat# 115-165-206, RRID:AB_2338695); anti-mouse IgG1 coupled to AlexaFluor647 (1/600; Thermo Fisher Scientific Cat# A-21240, RRID:AB_2535809); anti-rabbit IgG (H+L) coupled to AlexaFluor488 (1/400; Thermo Fisher Scientific Cat# A-21206, RRID:AB_2535792); anti-mouse IgG (H+L) coupled to AlexaFluor594 (1/200; Thermo Fisher Scientific Cat# A-21203, RRID:AB_2535789); anti-mouse IgG2a coupled to AlexaFluor488 (1/300; Thermo Fisher Scientific Cat# A-21131, RRID:AB_2535771); anti-mouse IgG1 coupled to AlexaFluor594 (1/300; Thermo Fisher Scientific Cat# A-21125, RRID:AB_2535767); anti-chicken IgY coupled to AlexaFluor594 (1/200; Jackson ImmunoResearch Labs Cat# 703-586-155, RRID:AB_2340378) and anti-rabbit IgG (H+L) coupled to AlexaFluor594 (1/400; Thermo Fisher Scientific Cat# A-21207, RRID:AB_141637).

### Confocal microscopy

Images were visualised using a Leica SP5 HyD resonant scanner Matrix screener inverted microscope equipped with a HCX PL APO CS 63x, 1.4 NA oil objective (Leica, Wetzlar, Germany). Triple or quadruple immunofluorescence was imaged with Leica Application Suite AF software (LAS AF; Leica Application Suite X, RRID:SCR_013673) after excitation of the Hoechst 33342 dye with a diode at a wavelength of 405 nm (452/75 Emission Filter), excitation of the AlexaFluor488 with an argon laser at a wavelength of 488 nm (525/50 Emission Filter), excitation of AlexaFluor594 with a diode DPSS at a wavelength of 561 nm (634/77 Emission Filter), excitation of Cy3 with a diode DPSS at a wavelength of 561 nm (595/49 Emission Filter), and excitation of AlexaFluor647 with a helium-neon laser at a wavelength of 633 nm (706/107 Emission Filter). Images were scanned sequentially to minimise cross excitation between channels and each line was scanned twice and averaged to increase the signal-to-noise ratio. The pinhole aperture was set to 1 airy. Images were acquired with 8x zoom at a resolution of 1024×1024. Z-stacks were acquired at 0.082 µm intervals, deconvolved and rendered using either Fiji (RRID:SCR_002285) or Icy (RRID:SCR_010587) software [80] (http://icy.bioimageanalysis.org/).

### Deconvolution of z-stacks and chromatic aberration correction

All confocal images were processed and analysed by using the Huygens Professional software version 19.04 (Scientific Volume Imaging, Huygens Software, RRID:SCR_014237). Deconvolution of confocal z-stacks was optimised using the following settings: automatic estimation of the average background with the mode “Lowest” and area radius = 0.7, deconvolution algorithm CMLE, maximum number of iterations = 40, signal to noise ratio (SNR) = 20, quality change threshold = 0.05, iteration mode = optimised, brick layout = automatic. Theoretical point spread function (PSF) values were estimated for each *z*-stack. All deconvolved images were corrected for chromatic shifts and for rotational differences between different channels using the Chromatic Aberration Corrector (CAC) from Huygens Professional software (Scientific Volume Imaging, Huygens Software, RRID:SCR_014237). To calibrate the image corrections, multifluorescent 0.2 µm TetraSpeck microspheres (ThermoFisher Scientific Cat#T7280) mounted on SlowFade Gold Antifade mountant (ThermoFisher Scientific Cat# S36937) were imaged with identical acquisition parameters. Images were deconvolved similarly, and were used to perform the chromatic aberration estimations with the cross correlation method in CAC software. Corrections were saved as templates and applied for correction of the similarly acquired and deconvolved images in CAC.

### Co-localisation analysis of confocal images

Co-localisation analysis was performed with the Co-localisation Analyzer plug-in of the Huygens Professional software (Scientific Volume Imaging, Huygens Software, RRID:SCR_014237, v19.04). Processed cross-section images (deconvolved and corrected for chromatic aberrations) of the parasites were opened with this plug-in and Pearson coefficients were calculated for each parasite. Specific areas of the parasite were cropped from the whole image (basal body area, flagella pocket area, mitotic spindle and reduced mitotic spindle areas, flagellar pocket neck area and flagellar tip area) and Pearson coefficients were calculated for these images. Pearson coefficients of the co-localisation in the basal body and flagellar pocket areas of (i) CK1.2-V5 with Centrin (CEN), IFT172, and DNA (Hoechst 33342, H); or (i) CEN with IFT172, from 14 images were plotted in scattered dot plots with the mean and standard deviation using GraphPad Prism 8.1.1 (GraphPad Software, GraphPad Prism, RRID:SCR_002798). Pearson coefficients of the co-localisation of CK1.2-V5 with tubulin in the mitotic spindle and reduced mitotic spindle areas from seven images were plotted similarly. Pearson coefficients of the co-localisation of CK1.2-V5 with Hsp90 from seven images and with Hsp70 from ten images were also plotted similarly.

### Epifluorescence microscopy and automated parasite detection

Images were visualised using a Zeiss upright widefield microscope equipped with Apotome2 grids and a Pln-Apo 63x, 1.4 NA oil objective (Zeiss). Light source used was a Mercury Lamp HXP 120, and following filters were used: DAPI (Excitation G365; dichroic FT 395; emission BP 420-470), FITC-A488-GFP (Excitation BP 455-495; dichroic FT 500; emission BP 505-555) and A594-TexasRed-mCherry-HcRed-mRFP (Excitation BP 542-582; dichroic FT 593; emission BP 604-644). Images were captured on an Axiocam MRm camera using ZEN Blue software. For comparison of different cell lines, identical parameters of acquisition were applied on all samples.

For the analysis of the fluorescence intensity in the parasite body of different cell lines (mock, WT and domain-deleted mutants), we used the graphical programming plugin Protocols in Icy software (Icy, RRID:SCR_010587 [81]). A screenshot of the protocol that was applied on the epifluorescence images is shown in Figure S6. Briefly, maximum intensity projection in Z was generated in all channels. Nuclei were segmented with HK-Means plugin (in the nucleus specific channel [82]), and the regions of interest (ROI) generated were used as input for automatically segment the boundary of the parasite body stained with V5 antibody (in all the cell lines) with Active Contours plugin [83]. The recovered ROIs were verified and corrected manually if needed. Properties of the ROI (e.g. sum fluorescence intensity, roundness, and interior) were obtained and used for analysis. Dot plots were generated with GraphPad Prism 8.1.1 (GraphPad Software, GraphPad Prism, RRID:SCR_002798).

### Protein extraction, SDS-PAGE and Western blot analysis

Logarithmic phase promastigotes were washed in DPBS and protein extraction was performed as described previously [45]. Ten micrograms of total protein were separated by SDS-PAGE, and transferred onto polyvinylidene difluoride (PVDF) membranes (Pierce). Membranes were blocked with 5% BSA in DPBS supplemented with 0.25% Tween20 (PBST) and incubated over night at 4°C with primary antibody mouse IgG2a anti-V5 tag monoclonal antibody (1/1000; Thermo Fisher Scientific Cat# R960-25, RRID:AB_2556564) in 2,5% BSA in PBST. Membranes were then washed in PBST and incubated with secondary antibody anti-mouse IgG (H+L) coupled to horseradish peroxidase (1/20000; ThermoFisher Scientific Cat# 32230, RRID:AB_1965958). Proteins were revealed by SuperSignal™ West Pico Chemiluminescent Substrate (ThermoFisher Scientific Cat# 34580) using the PXi image analysis system (Syngene) at various exposure times. Membranes were then stained with Bio-Safe Coomassie (Bio-Rad Cat #1610786) to serve as loading controls.

### Proteasome and lysosome inhibition assays

Logarithmic phase promastigotes expressing CK1.2-V5-His_6_, CK1.2ΔC10-V5-His_6_, CK1.2ΔC43-V5- His_6_, CK1.2ΔN7-V5-His_6_ or mock control (pLEXSY empty plasmid) were resuspended into fresh M199-supplemented promastigote medium at 5×10^6^ parasites per mL with or without either 10 µM MG132 (Sigma-Aldrich Cat# M7449) or 20 mM NH_4_Cl (VWR Chemicals Cat# 21235.297). Drug selection was maintained with 30 µg hygromycin B (Invitrogen). Parasites were grown for 24h at 26°C and were then lysed for protein extraction as described before. Western blot analysis of ten micrograms of total protein was performed as described before.

Ten micrograms of total proteins treated with or without MG132 were also subjected to Western blot analysis to detect ubiquitinylated proteins. Membrane was blocked with 5% BSA in DPBS supplemented with 0.25% Tween20 (PBST) and incubated over night at 4°C with primary mouse mono- and polyubiquitinylated conjugates FK2 monoclonal antibody (1/500; Enzo Life Sciences Cat# BML-PW8810, RRID:AB_10541840) in 2,5% BSA in PBST. Following washing in PBST, the membrane was incubated with secondary antibody anti-mouse IgG (H+L) coupled to horseradish peroxidase (1/20000; ThermoFisher Scientific Cat# 32230, RRID:AB_1965958). The immunoblot was revealed with SuperSignal™ West Pico PLUS Chemiluminescent Substrate (ThermoFisher Scientific Cat# 34580) using the PXi image analysis system (Syngene) with 5 min exposure time. The membrane was then stained with Bio-Safe Coomassie G-250 stain (Bio-Rad Cat #1610786) to serve as loading control.

To validate lysosomal inhibition by NH_4_Cl treatment, parasites were sampled prior cell lysis and stained to access lysosomal pH. Treated or untreated parasites were incubated with 100 mM LysoTracker™ Red DND-99 (ThermoFisher Scientific Cat# L7528) in culture medium for 30 min at 26°C and analysed with a CytoFLEX flow cytometer (Beckman Coulter, Inc.) to test for acidic pH of lysosomes upon treatment (exλ = 577 nm; emλ = 590 nm). Lysotracker fluorescence intensity was measured for 15000 parasites using CytExpert software (CytExpert Software, RRID:SCR_017217, Beckman Coulter, v2.2.0.97). Graphs representing mean Lysotracker fluorescence intensity were generated with GraphPad Prism 8.1.1 (GraphPad Software, GraphPad Prism, RRID:SCR_002798).

### Recombinant expression, purification of CK1.2-V5-His_6_, CK1.2ΔC10-V5-His_6_, CK1.2ΔC43-V5-His_6_ and CK1.2ΔN7-V5-His_6_ and protein kinase assay

*Escherichia coli* Rosetta (DE3) pLysS Competent Cells (Merck Cat# 70956) containing pBAD-thio-topo-LmaCK1.2-V5-His_6_, pBAD-thio-topo-LmaCK1.2ΔC10-V5-His_6_, pBAD-thio-topo-LmaCK1.2ΔC43-V5-His_6_ or pBAD-thio-topo-LmaCK1.2ΔN7-V5-His6 were grown at 37°C and induced with arabinose (0,02% final) for 4h at room temperature [13]. Cells were harvested by centrifugation at 10,000 g for 10 min at 4°C and the recombinant proteins were purified as described previously [13] [73]. The eluates were supplemented with 15% glycerol and stored at −80°C. The kinase assays were performed as described previously [13] [73].

## QUANTIFICATION AND STATISTICAL ANALYSIS

Statistical analyses were performed with GraphPad Prism 8.1.1 (GraphPad Software, GraphPad Prism, RRID:SCR_002798) using unpaired t test (parametric test). Graphs were drawn using the same software. All errors correspond to the 95% confidence interval. Statistically significant differences are indicated with three (p<0.01), four (p<0.001) or five asterisks (p<0.0001). The number of samples analysed for each experiment is indicated in figure legends.

## RESOURCES TABLE

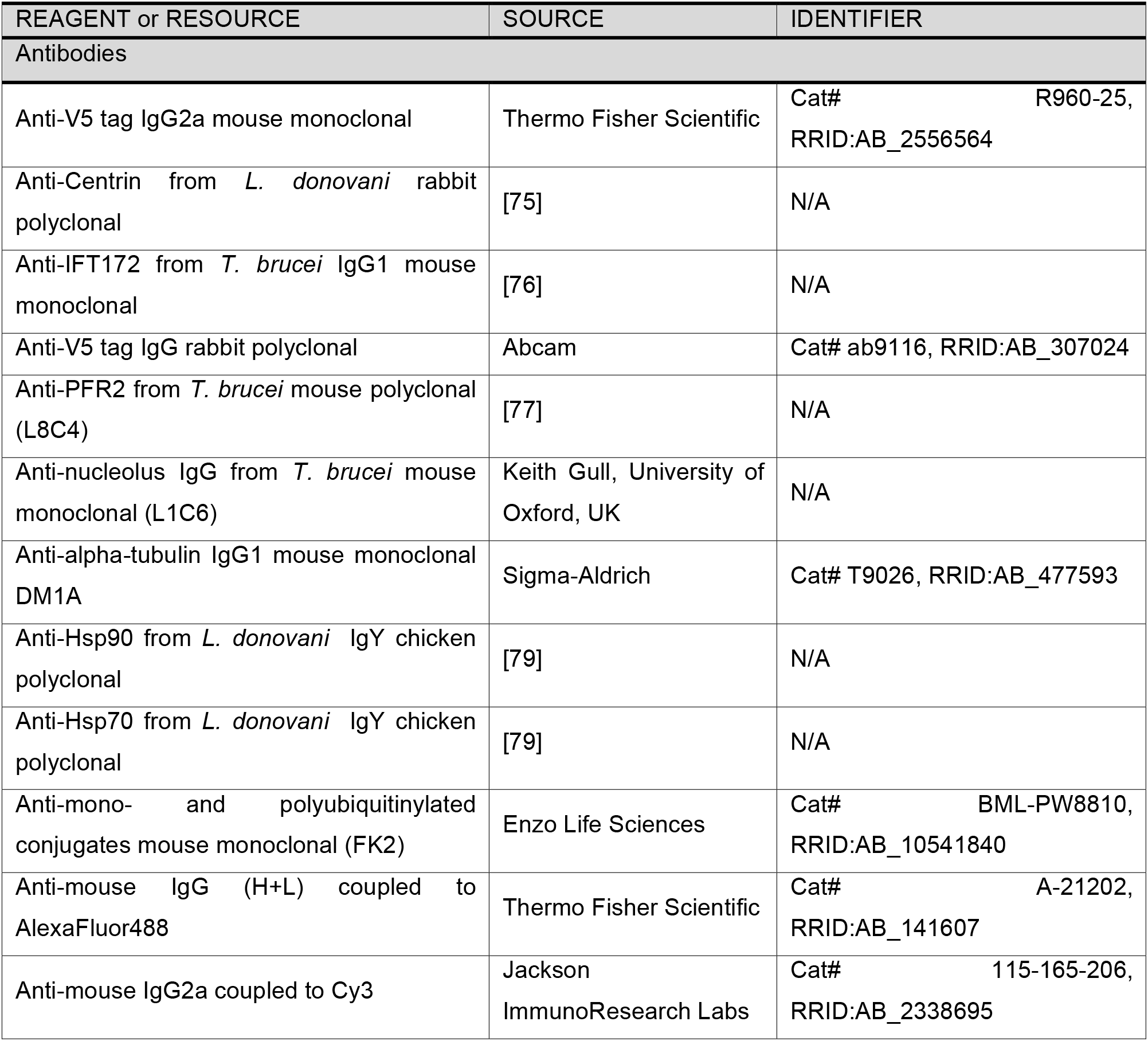

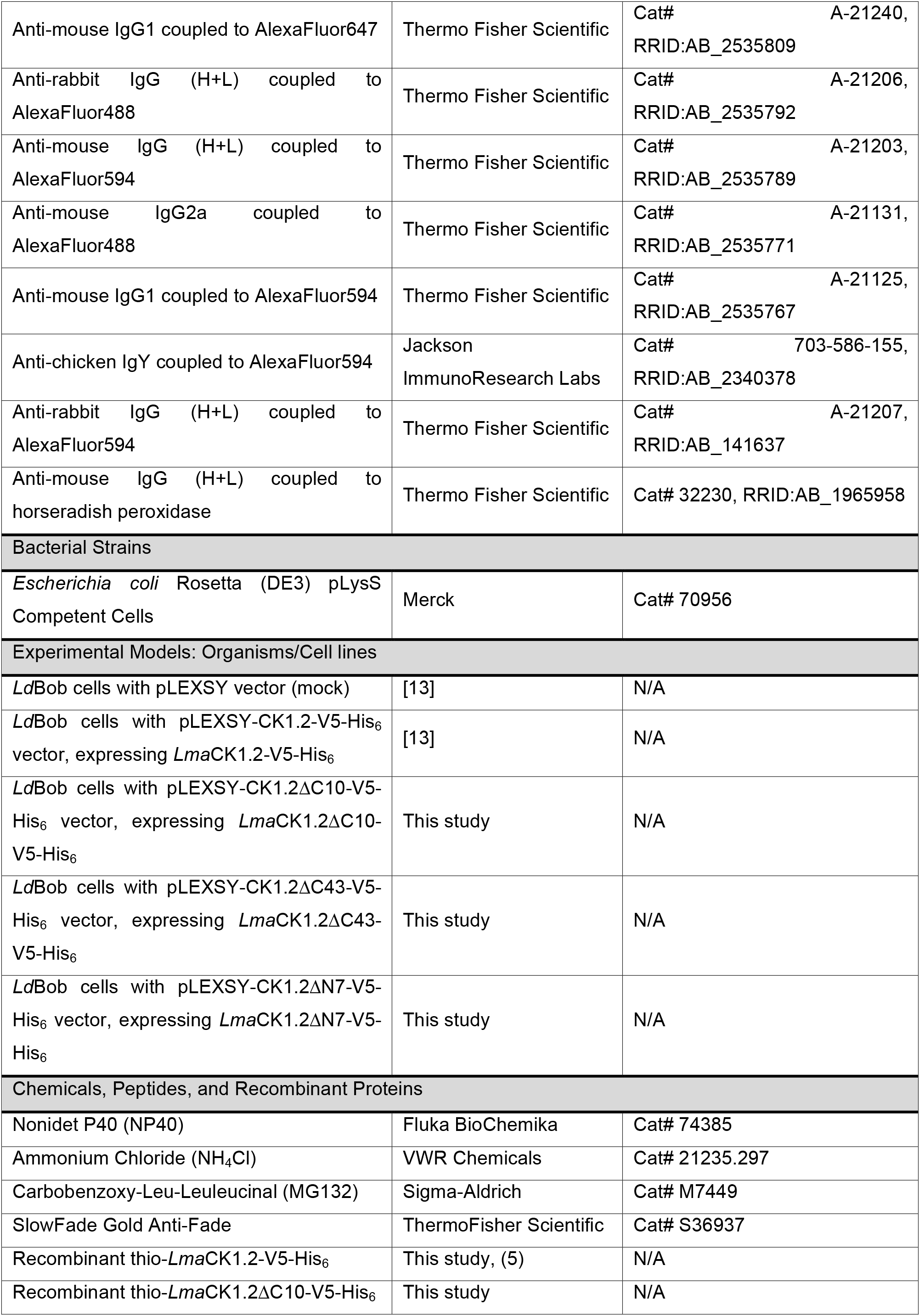

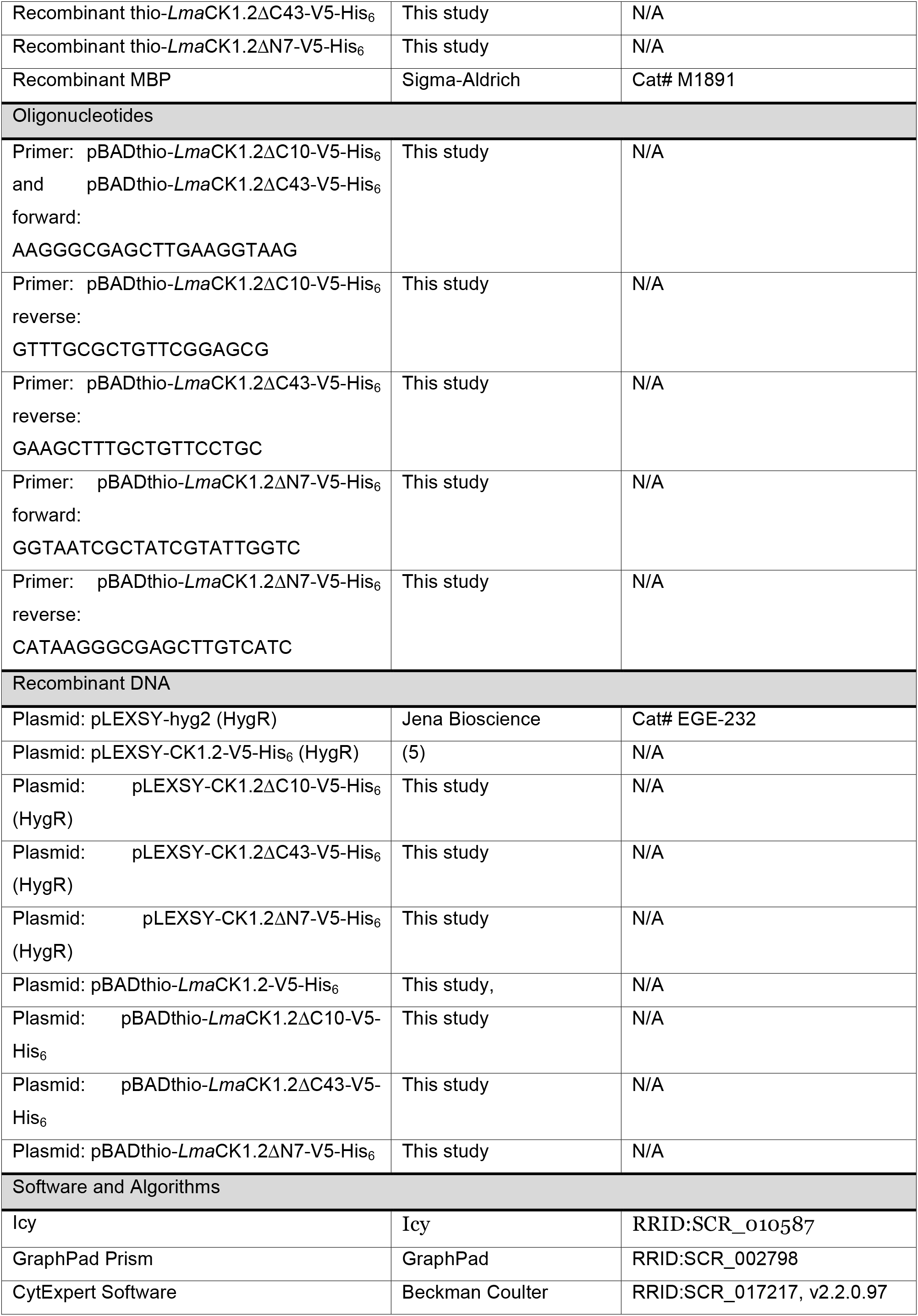

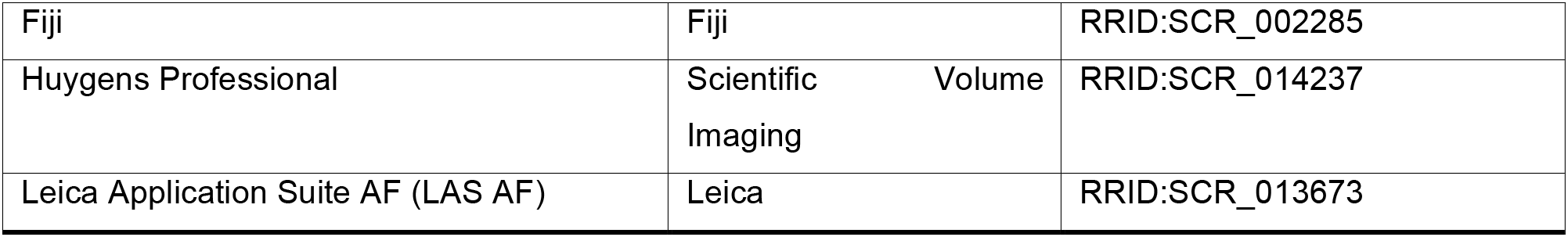

## Acknowledgments

This work was supported by the ANR-13-ISV3-0009. Daniel Martel was supported by the French Government’s Investissements d’Avenir program Laboratoire d’Excellence Integrative Biology of Emerging Infectious Diseases (grant no. ANR-10-LABX-62-IBEID studentship). Part of the work was supported by the Deutsche Forschungsgemeinschaft Grant Cl 120/8-1. We thank Frauke Fuchs for technical assistance. The authors would like to thanks Hira L Nakhasi, U.S. FDA, for the anti-*Ld*centrin antibody; Philippe Bastin, Institut Pasteur, for the anti-*Tb*IFT172 and anti-*Tb*PFR2 (L8C4) antibodies; Keith Gull, University of Oxford for the L1C6 antibody; the Unit of Technology and Service - Photonic BioImaging (UTechS PBI) from the Institut Pasteur, for the help with confocal microscopy and analyses of co-localisations, in particular Audrey Salles, Julien Fernandes and Anne Danckaert. Finally, we would like to thanks Brice Rotureau, Thierry Blisnick and Philippe Bastin for fruitful discussions and advices.

## Author contribution

Conceptualization: DM, NR; Formal analysis: DM, JC, GFS, NR; Investigation: DM, SP, KB, NR; Funding acquisition: JC, GFS, NR; Supervision JC, GFS, NR; Writing –original draft: DM, NR; Writing –review & Editing: DM, SP, KB, JC, GFS, NR.

## Supplemental figure legends

**Figure S1: CK1.2-V5 localisation in methanol-fixed promastigotes.**

IFA of *Ld*Bob pLEXSY-CK1.2-V5 (A) and *Ld*Bob pLEXSY (mock, B) promastigotes, fixed in ice-cold methanol for 3 minutes and stained with anti-V5 antibody to detect CK1.2-V5 localisation. The epifluorescence images were acquired under the same conditions and show the anti-V5 staining (CK1.2-V5 or V5), Hoechst 33342 staining (H), a merge of the anti-V5 (green) and H (red) signals and the transmission image (Trans). Scale bar, 5 µm. The pictures are maximum intensity projection of the z-stacks containing the parasites.

**Figure S2: Hsp90 and Hsp70 cytoplasmic localisation.**

(A) IFA of *Ld*Bob pLEXSY-CK1.2-V5 promastigotes fixed with PFA, and stained with anti-*Ld*Hsp90 antibody to detect Hsp90 localisation. The single channel images show Hsp90 and Hoechst 33342 (H) signals and the transmission image (Trans). The merged channel shows Hsp90 (red) and H (blue) signals. Scale bar, 2 µm. These pictures are single stacks extracted from deconvolved confocal stacks corrected for chromatic aberration. (B) IFA of *Ld*Bob pLEXSY-CK1.2-V5 promastigotes fixed with PFA without detergent treatment, and stained with anti-Hsp70 antibody. The single channel images show Hsp70 and Hoechst 33342 (H) signals and the transmission image (Trans). The merged channel shows Hsp70 (red) and H (blue) signals. Scale bar, 2 µm. These pictures are single stacks extracted from deconvolved confocal stacks corrected for chromatic aberration.

**Figure S3: CK1.2 localises in the nucleolus and to the mitotic spindle.**

(A) IFA pictures of *Ld*Bob pLEXSY-CK1.2-V5 promastigotes obtained after detergent treatment followed by PFA fixation and stained with anti-V5 (CK1.2-V5) and anti-L1C6 (nucleolus, L1C6) antibodies. Confocal images representing sequential events of mitosis revealed different localisation patterns of L1C6 nucleolar marker and CK1.2-V5. (a – f) The images correspond to the transmission (Trans), the merged containing CK1.2-V5 (green), Hoechst 33342 (H) (blue) and L1C6 (red) signals. The following four images show a magnification of the nuclear region with the merged and single channel images. N=nucleus, K=kinetoplast. Scale bar, 2 µm or 1 µm for magnified images. These pictures are single stacks extracted from deconvolved confocal stacks corrected for chromatic aberration. (B) IFA pictures of *Ld*Bob pLEXSY-CK1.2-V5 promastigotes obtained after detergent treatment followed by PFA fixation and stained with anti-V5 and anti-α-tubulin antibodies. Sequential images of various stages of cell division (a – e) showing the single channel images for CK1.2-V5, H and α-tubulin signals, the merged images showing CK1.2-V5 (green), H (blue) and α-tubulin (red) signals, and the transmission image (Trans). Scale bar, 2 µm. These pictures are single stacks extracted from deconvolved confocal stacks corrected for chromatic aberration.

**Figure S4: Inhibition of proteasomal and lysosomal degradation.**

(A) Proteosomal degradation. (a) Logarithmic phase promastigotes, from the same cell lines as in (Figure 6C), were treated by the proteasome inhibitor MG132 for 18h. Proteins from treated and untreated control were extracted and twenty micrograms were analysed by Western blotting (WB) using α-V5 (top panel). The Coomassie-stained membrane of the blot is included as a loading control (bottom panels). MW= Molecular Weight. The blots are representative of three independent experiments. (b) Logarithmic phase promastigotes from *Ld*Bob pLEXSY-CK1.2-V5 (WT) or expressing truncated kinase mutants *Ld*Bob pLEXSY-CK1.2-ΔC10-V5 (ΔC10), *Ld*Bob pLEXSY-CK1.2-ΔC43-V5 (ΔC43) and *Ld*Bob pLEXSY-CK1.2-ΔN7-V5 (ΔN7) were treated with the proteasome inhibitor MG132 for 18h. Proteins from treated and untreated samples were extracted and twenty micrograms were analysed by Western blotting using the mono- and poly-ubiquitinylated conjugates monoclonal (FK2) antibody (α-Ubiquitin) (left panel). The Coomassie-stained membrane of the blot is included as a loading control (right panel). MW= Molecular Weight. The blot is representative of two independent experiments. (B) Lysosomal degradation. (a) Logarithmic phase promastigotes from the same cell lines as in (Figure 6C) were treated by ammonium chloride (NH_4_Cl), an inhibitor of lysosomal degradation, for 18h. Proteins from treated and untreated control were extracted and twenty micrograms were analysed by Western blotting (WB) using α-V5 (top panel). The Coomassie-stained membrane of the blot is included as a loading control (bottom panels). MW= Molecular Weight. The blots are representative of three independent experiments. (b) Logarithmic phase promastigotes from *Ld*Bob pLEXSY-CK1.2-V5 (WT) or expressing truncated kinase mutants *Ld*Bob pLEXSY-CK1.2-ΔC10-V5 (ΔC10), *Ld*Bob pLEXSY-CK1.2-ΔC43-V5 (ΔC43) and *Ld*Bob pLEXSY-CK1.2-ΔN7-V5 (ΔN7) were treated with ammonium chloride for 18h to inhibit the lysosomal degradation. Untreated parasites were used as control. Lysosomes of the parasites were then stained with 100 mM LysoTracker™ Red DND-99 for 30 min at 26°C. Mean fluorescence intensity of the Lysotracker accumulation in lysosomes was measured by flow cytometry in treated (black) and untreated (grey) parasites for ∼15000 parasites. The data are representative of two independent experiments. The graph was generated with GraphPad Prism software.

**Figure S5: Position of the LCR in human CK1s.**

Cartoon representing the low complexity region (LCR, dark grey) on the protein sequence (light grey) of human CK1. CK1A (P48729), CK1D (P48730), CK1E (P49674), CK1G1 (Q9HCP0), CK1G2 (P78368) and CK1G3 (Q9Y6M4).

**Figure S6: Protocol for automatic segmentation of parasite bodies.**

Screenshot of the protocol applied on the epifluorescence images to analyse diverse parameters of the parasite body of *Ld*Bob pLEXSY-CK1.2-V5, mock or domain-deleted mutants, stained with the anti- V5 antibody to detect CK1.2-V5 localisation. Protocol is a graphical programming plugin in Icy software (Icy, RRID:SCR010)

## References

1. Knippschild, U., Kruger, M., Richter, J., Xu, P., Garcia-Reyes, B., Peifer, C., Halekotte, J., Bakulev, V., and Bischof, J. (2014). The CK1 Family: Contribution to Cellular Stress Response and Its Role in Carcinogenesis. Front Oncol 4, 96.

2. Knippschild, U., Gocht, A., Wolff, S., Huber, N., Lohler, J., and Stoter, M. (2005). The casein kinase 1 family: participation in multiple cellular processes in eukaryotes. Cell Signal 17, 675–689.

3. Schittek, B., and Sinnberg, T. (2014). Biological functions of casein kinase 1 isoforms and putative roles in tumorigenesis. Mol Cancer 13, 231.

4. Jiang, S., Zhang, M., Sun, J., and Yang, X. (2018). Casein kinase 1alpha: biological mechanisms and theranostic potential. Cell Commun Signal 16, 23.

5. Greer, Y.E., Westlake, C.J., Gao, B., Bharti, K., Shiba, Y., Xavier, C.P., Pazour, G.J., Yang, Y., and Rubin, J.S. (2014). Casein kinase 1delta functions at the centrosome and Golgi to promote ciliogenesis. Molecular biology of the cell 25, 1629–1640.

6. Greer, Y.E., and Rubin, J.S. (2011). Casein kinase 1 delta functions at the centrosome to mediate Wnt-3a-dependent neurite outgrowth. J Cell Biol 192, 993–1004.

7. Peng, Y., Moritz, M., Han, X., Giddings, T.H., Lyon, A., Kollman, J., Winey, M., Yates, J., 3rd, Agard, D.A., Drubin, D.G., et al. (2015b). Interaction of CK1delta with gammaTuSC ensures proper microtubule assembly and spindle positioning. Molecular biology of the cell 26, 2505-2518.

8. Kafadar, K.A., Zhu, H., Snyder, M., and Cyert, M.S. (2003). Negative regulation of calcineurin signaling by Hrr25p, a yeast homolog of casein kinase I. Genes Dev 17, 2698–2708.

9. Lusk, C.P., Waller, D.D., Makhnevych, T., Dienemann, A., Whiteway, M., Thomas, D.Y., and Wozniak, R.W. (2007). Nup53p is a target of two mitotic kinases, Cdk1p and Hrr25p. Traffic 8, 647-660.

10. Peng, Y., Grassart, A., Lu, R., Wong, C.C.L., Yates, J., 3rd, Barnes, G., and Drubin, D.G. (2015a). Casein kinase 1 promotes initiation of clathrin-mediated endocytosis. Dev Cell 32, 231-240.

11. Zhang, B., Shi, Q., Varia, S.N., Xing, S., Klett, B.M., Cook, L.A., and Herman, P.K. (2016). The Activity-Dependent Regulation of Protein Kinase Stability by the Localization to P-Bodies. Genetics 203, 1191–1202.

12. Dorin-Semblat, D., Demarta-Gatsi, C., Hamelin, R., Armand, F., Carvalho, T.G., Moniatte, M., and Doerig, C. (2015). Malaria Parasite-Infected Erythrocytes Secrete PfCK1, the Plasmodium Homologue of the Pleiotropic Protein Kinase Casein Kinase 1. PloS one 10, e0139591.

13. Rachidi, N., Taly, J.F., Durieu, E., Leclercq, O., Aulner, N., Prina, E., Pescher, P., Notredame, C., Meijer, L., and Spath, G.F. (2014). Pharmacological assessment defines the Leishmania donovani casein kinase 1 as a drug target and reveals important functions in parasite viability and intracellular infection. Antimicrob Agents Chemother 58.

14. Silverman, J.M., and Reiner, N.E. (2010). Exosomes and other microvesicles in infection biology: organelles with unanticipated phenotypes. Cell Microbiol 13, 1–9.

15. Jayaswal, S., Kamal, M.A., Dua, R., Gupta, S., Majumdar, T., Das, G., Kumar, D., and Rao, K.V. (2010). Identification of host-dependent survival factors for intracellular Mycobacterium tuberculosis through an siRNA screen. PLoS pathogens 6, e1000839.

16. Xia, C., Wolf, J.J., Vijayan, M., Studstill, C.J., Ma, W., and Hahm, B. (2018). Casein Kinase 1alpha Mediates the Degradation of Receptors for Type I and Type II Interferons Caused by Hemagglutinin of Influenza A Virus. J Virol 92.

17. Zhang, L., Li, H., Chen, Y., Gao, X., Lu, Z., Gao, L., Wang, Y., Gao, Y., Gao, H., Liu, C., et al. (2017). The down-regulation of casein kinase 1 alpha as a host defense response against infectious bursal disease virus infection. Virology 512, 211–221.

18. Cegielska, A., and Virshup, D.M. (1993). Control of simian virus 40 DNA replication by the HeLa cell nuclear kinase casein kinase I. Mol Cell Biol 13, 1202–1211.

19. Sudha, G., Yamunadevi, S., Tyagi, N., Das, S., and Srinivasan, N. (2012). Structural and molecular basis of interaction of HCV non-structural protein 5A with human casein kinase 1alpha and PKR. BMC structural biology 12, 28.

20. Bhattacharya, D., Ansari, I.H., and Striker, R. (2009). The flaviviral methyltransferase is a substrate of Casein Kinase 1. Virus research 141, 101–104.

21. Solyakov, L., Halbert, J., Alam, M.M., Semblat, J.P., Dorin-Semblat, D., Reininger, L., Bottrill, A.R., Mistry, S., Abdi, A., Fennell, C., et al. (2011). Global kinomic and phospho-proteomic analyses of the human malaria parasite Plasmodium falciparum. Nat Commun 2, 565.

22. Silverman, J.M., Clos, J., de’Oliveira, C.C., Shirvani, O., Fang, Y., Wang, C., Foster, L.J., and Reiner, N.E. (2010). An exosome-based secretion pathway is responsible for protein export from Leishmania and communication with macrophages. J Cell Sci 123, 842–852.

23. Dan-Goor, M., Nasereddin, A., Jaber, H., and Jaffe, C.L. (2013). Identification of a secreted casein kinase 1 in Leishmania donovani: effect of protein over expression on parasite growth and virulence. PloS one 8, e79287.

24. Atayde, V.D., Suau, H.A., Townsend, S., Hassani, K., Kamhawi, S., and Olivier, M. (2015). Exosome secretion by the parasitic protozoan Leishmania within the sand fly midgut. Cell reports 13, 957–967.

25. Liu, J., Carvalho, L.P., Bhattacharya, S., Carbone, C.J., Kumar, K.G., Leu, N.A., Yau, P.M., Donald, R.G., Weiss, M.J., Baker, D.P., et al. (2009). Mammalian casein kinase 1alpha and its leishmanial ortholog regulate stability of IFNAR1 and type I interferon signaling. Mol Cell Biol 29, 6401–6412.

26. Xu, P., Ianes, C., Gartner, F., Liu, C., Burster, T., Bakulev, V., Rachidi, N., Knippschild, U., and Bischof, J. (2019). Structure, regulation, and (patho-)physiological functions of the stress-induced protein kinase CK1 delta (CSNK1D). Gene 715, 144005.

27. Fulcher, L.J., Bozatzi, P., Tachie-Menson, T., Wu, K.Z.L., Cummins, T.D., Bufton, J.C., Pinkas, D.M., Dunbar, K., Shrestha, S., Wood, N.T., et al. (2018). The DUF1669 domain of FAM83 family proteins anchor casein kinase 1 isoforms. Science signaling 11.

28. Vaughan, S., and Gull, K. (2015). Basal body structure and cell cycle-dependent biogenesis in Trypanosoma brucei. Cilia 5, 5.

29. Selvapandiyan, A., Kumar, P., Morris, J.C., Salisbury, J.L., Wang, C.C., Nakhasi, H.L., and Cohen-Fix, O. (2007). Centrin1 Is Required for Organelle Segregation and Cytokinesis in Trypanosoma brucei. Molecular Biology of the Cell 18, 3290–3301.

30. Shi, J., Franklin, J.B., Yelinek, J.T., Ebersberger, I., Warren, G., and He, C.Y. (2008). Centrin4 coordinates cell and nuclear division in T. brucei. Journal of cell science 121, 3062–3070.

31. Sunter, J.D., Yanase, R., Wang, Z., Catta-Preta, C.M.C., Moreira-Leite, F., Myskova, J., Pruzinova, K., Volf, P., Mottram, J.C., and Gull, K. (2019). Leishmania flagellum attachment zone is critical for flagellar pocket shape, development in the sand fly, and pathogenicity in the host. Proceedings of the National Academy of Sciences of the United States of America 116, 6351–6360.

32. Eliaz, D., Kannan, S., Shaked, H., Arvatz, G., Tkacz, I.D., Binder, L., Waldman Ben-Asher, H., Okalang, U., Chikne, V., Cohen-Chalamish, S., et al. (2017). Exosome secretion affects social motility in Trypanosoma brucei. PLoS pathogens 13, e1006245.

33. Thery, C., Ostrowski, M., and Segura, E. (2009). Membrane vesicles as conveyors of immune responses. Nat Rev Immunol 9, 581–593.

34. Muller, P., Ruckova, E., Halada, P., Coates, P.J., Hrstka, R., Lane, D.P., and Vojtesek, B. (2012). C-terminal phosphorylation of Hsp70 and Hsp90 regulates alternate binding to co-chaperones CHIP and HOP to determine cellular protein folding/degradation balances. Oncogene.

35. Hombach-Barrigah, A., Bartsch, K., Smirlis, D., Rosenqvist, H., MacDonald, A., Dingli, F., Loew, D., Spath, G.F., Rachidi, N., Wiese, M., et al. (2019). Leishmania donovani 90 kD Heat Shock Protein - Impact of Phosphosites on Parasite Fitness, Infectivity and Casein Kinase Affinity. Scientific reports 9, 5074.

36. Demmel, L., Schmidt, K., Lucast, L., Havlicek, K., Zankel, A., Koestler, T., Reithofer, V., de Camilli, P., and Warren, G. (2016). The endocytic activity of the flagellar pocket in Trypanosoma brucei is regulated by an adjacent phosphatidylinositol phosphate kinase. J Cell Sci 129, 2285.

37. Wheeler, R.J., Sunter, J.D., and Gull, K. (2016). Flagellar pocket restructuring through the Leishmania life cycle involves a discrete flagellum attachment zone. J Cell Sci 129, 854–867.

38. Názer, E., and Sánchez, D.O. (2011). Nucleolar Accumulation of RNA Binding Proteins Induced by ActinomycinD Is Functional in Trypanosoma cruzi and Leishmania mexicana but Not in T. brucei. PloS one 6, e24184.

39. Motta, M.C.M., Souza, W.d., and Thiry, M. (2003). Immunocytochemical detection of DNA and RNA in endosymbiont-bearing trypanosomatids. FEMS Microbiology Letters 221, 17–23.

40. Ogbadoyi, E., Ersfeld, K., Robinson, D., Sherwin, T., and Gull, K. (2000). Architecture of the Trypanosoma brucei nucleus during interphase and mitosis. Chromosoma 108, 501–513.

41. Kumar, G., Kajuluri, L.P., Gupta, C.M., and Sahasrabuddhe, A.A. (2016). A twinfilin-like protein coordinates karyokinesis by influencing mitotic spindle elongation and DNA replication in Leishmania. Mol Microbiol 100, 173–187.

42. Stoter, M., Bamberger, A.-M., Aslan, B., Kurth, M., Speidel, D., Loning, T., Frank, H.-G., Kaufmann, P., Lohler, J., Henne-Bruns, D., et al. (2005). Inhibition of casein kinase I delta alters mitotic spindle formation and induces apoptosis in trophoblast cells. Oncogene 24, 7964–7975.

43. Boucher, N., Dacheux, D., Giroud, C., and Baltz, T. (2007). An essential cell cycle-regulated nucleolar protein relocates to the mitotic spindle where it is involved in mitotic progression in Trypanosoma brucei. J Biol Chem 282, 13780–13790.

44. Schittek, B., and Sinnberg, T. (2014). Biological functions of casein kinase 1 isoforms and putative roles in tumorigenesis. Mol Cancer 13, 231.

45. Martel, D., Beneke, T., Gluenz, E., Spath, G.F., and Rachidi, N. (2017). Characterisation of Casein Kinase 1.1 in Leishmania donovani Using the CRISPR Cas9 Toolkit. BioMed research international 2017, 4635605.

46. Qin, H., Shao, Q., Igdoura, S.A., Alaoui-Jamali, M.A., and Laird, D.W. (2003). Lysosomal and proteasomal degradation play distinct roles in the life cycle of Cx43 in gap junctional intercellular communication-deficient and -competent breast tumor cells. J Biol Chem 278, 30005–30014.

47. Besteiro, S. (2017). Autophagy in apicomplexan parasites. Current opinion in microbiology 40, 14–20.

48. Crawford, L.J., Walker, B., Ovaa, H., Chauhan, D., Anderson, K.C., Morris, T.C., and Irvine, A.E. (2006). Comparative selectivity and specificity of the proteasome inhibitors BzLLLCOCHO, PS-341, and MG-132. Cancer Res 66, 6379-6386.

49. Donald, R.G., Zhong, T., Meijer, L., and Liberator, P.A. (2005). Characterization of two T. gondii CK1 isoforms. Mol Biochem Parasitol 141, 15–27.

50. Hubstenberger, A., Courel, M., Benard, M., Souquere, S., Ernoult-Lange, M., Chouaib, R., Yi, Z., Morlot, J.B., Munier, A., Fradet, M., et al. (2017). P-Body Purification Reveals the Condensation of Repressed mRNA Regulons. Mol Cell 68, 144–157 e145.

51. Dallagiovanna, B., Correa, A., Probst, C.M., Holetz, F., Smircich, P., de Aguiar, A.M., Mansur, F., da Silva, C.V., Mortara, R.A., Garat, B., et al. (2008). Functional genomic characterization of mRNAs associated with TcPUF6, a pumilio-like protein from Trypanosoma cruzi. J Biol Chem 283, 8266–8273.

52. Kramer, S., Queiroz, R., Ellis, L., Webb, H., Hoheisel, J.D., Clayton, C., and Carrington, M. (2008). Heat shock causes a decrease in polysomes and the appearance of stress granules in trypanosomes independently of eIF2(alpha) phosphorylation at Thr169. J Cell Sci 121, 3002–3014.

53. Cassola, A., De Gaudenzi, J.G., and Frasch, A.C. (2007). Recruitment of mRNAs to cytoplasmic ribonucleoprotein granules in trypanosomes. Mol Microbiol 65, 655–670.

54. Minia, I., and Clayton, C. (2016). Regulating a Post-Transcriptional Regulator: Protein Phosphorylation, Degradation and Translational Blockage in Control of the Trypanosome Stress-Response RNA-Binding Protein ZC3H11. PLoS pathogens 12, e1005514.

55. Droll, D., Minia, I., Fadda, A., Singh, A., Stewart, M., Queiroz, R., and Clayton, C. (2013). Post-transcriptional regulation of the trypanosome heat shock response by a zinc finger protein. PLoS pathogens 9, e1003286.

56. Lee, K.H., Johmura, Y., Yu, L.R., Park, J.E., Gao, Y., Bang, J.K., Zhou, M., Veenstra, T.D., Yeon Kim, B., and Lee, K.S. (2012). Identification of a novel Wnt5a-CK1varepsilon-Dvl2-Plk1-mediated primary cilia disassembly pathway. The EMBO journal 31, 3104–3117.

57. Subota, I., Julkowska, D., Vincensini, L., Reeg, N., Buisson, J., Blisnick, T., Huet, D., Perrot, S., Santi-Rocca, J., Duchateau, M., et al. (2014). Proteomic analysis of intact flagella of procyclic Trypanosoma brucei cells identifies novel flagellar proteins with unique sub-localization and dynamics. Molecular & cellular proteomics: MCP 13, 1769–1786.

58. Beneke, T., Demay, F., Hookway, E., Ashman, N., Jeffery, H., Smith, J., Valli, J., Becvar, T., Myskova, J., Lestinova, T., et al. (2018). Genetic dissection of a *Leishmania* flagellar proteome demonstrates requirement for directional motility in sand fly infections. bioRxiv, 476994.

59. Boesger, J., Wagner, V., Weisheit, W., and Mittag, M. (2012). Application of phosphoproteomics to find targets of casein kinase 1 in the flagellum of chlamydomonas. International journal of plant genomics 2012, 581460.

60. Perry, J.A., Sinclair-Davis, A.N., McAllaster, M.R., and de Graffenried, C.L. (2018). TbSmee1 regulates hook complex morphology and the rate of flagellar pocket uptake in Trypanosoma brucei. Mol Microbiol 107, 344–362.

61. Zemp, I., Wandrey, F., Rao, S., Ashiono, C., Wyler, E., Montellese, C., and Kutay, U. (2014). CK1delta and CK1epsilon are components of human 40S subunit precursors required for cytoplasmic 40S maturation. J Cell Sci 127, 1242–1253.

62. Ghalei, H., Schaub, F.X., Doherty, J.R., Noguchi, Y., Roush, W.R., Cleveland, J.L., Stroupe, M.E., and Karbstein, K. (2015). Hrr25/CK1delta-directed release of Ltv1 from pre-40S ribosomes is necessary for ribosome assembly and cell growth. J Cell Biol 208, 745–759.

63. Andersen, J.S., Lam, Y.W., Leung, A.K., Ong, S.E., Lyon, C.E., Lamond, A.I., and Mann, M. (2005). Nucleolar proteome dynamics. Nature 433, 77–83.

64. Zhou, Q., Lee, K.J., Kurasawa, Y., Hu, H., An, T., and Li, Z. (2018). Faithful chromosome segregation in Trypanosoma brucei requires a cohort of divergent spindle-associated proteins with distinct functions. Nucleic acids research 46, 8216–8231.

65. Fulcher, L.J., He, Z., Mei, L., Macartney, T., Wood, N., Prescott, A.R., Whigham, A., Varghese, J., Gourlay, R., Ball, G., et al. (2018). FAM83D directs protein kinase CK1α to the mitotic spindle for proper spindle positioning. bioRxiv, 480616.

66. Urbaniak, M.D. (2009). Casein kinase 1 isoform 2 is essential for bloodstream form Trypanosoma brucei. Molecular and Biochemical Parasitology 166, 183–185.

67. Coletta, A., Pinney, J.W., Solis, D.Y., Marsh, J., Pettifer, S.R., and Attwood, T.K. (2010). Low-complexity regions within protein sequences have position-dependent roles. BMC systems biology 4, 43.

68. Aslett, M., Aurrecoechea, C., Berriman, M., Brestelli, J., Brunk, B.P., Carrington, M., Depledge, D.P., Fischer, S., Gajria, B., Gao, X., et al. (2010). TriTrypDB: a functional genomic resource for the Trypanosomatidae. Nucleic acids research 38, D457–462.

69. Silverman, J.M., Clos, J., Horakova, E., Wang, A.Y., Wiesgigl, M., Kelly, I., Lynn, M.A., McMaster, W.R., Foster, L.J., Levings, M.K., et al. (2010). Leishmania exosomes modulate innate and adaptive immune responses through effects on monocytes and dendritic cells. J Immunol 185, 5011–5022.

70. Schultz, J., Milpetz, F., Bork, P., and Ponting, C.P. (1998). SMART, a simple modular architecture research tool: identification of signaling domains. Proceedings of the National Academy of Sciences of the United States of America 95, 5857–5864.

71. Sacerdoti-Sierra, N., and Jaffe, C.L. (1997). Release of ecto-protein kinases by the protozoan parasite Leishmania major. J Biol Chem 272, 30760–30765.

72. Bohm, T., Meng, Z., Haas, P., Henne-Bruns, D., Rachidi, N., Knippschild, U., and Bischof, J. (2019). The kinase domain of CK1delta can be phosphorylated by Chk1. Bioscience, biotechnology, and biochemistry, 1-13.

73. Durieu, E., Prina, E., Leclercq, O., Oumata, N., Gaboriaud-Kolar, N., Vougogiannopoulou, K., Aulner, N., Defontaine, A., No, J.H., Ruchaud, S., et al. (2016). From Drug Screening to Target Deconvolution: a Target-Based Drug Discovery Pipeline Using Leishmania Casein Kinase 1 Isoform 2 To Identify Compounds with Antileishmanial Activity. Antimicrobial Agents and Chemotherapy 60, 2822–2833.

74. Florimond, C., Sahin, A., Vidilaseris, K., Dong, G., Landrein, N., Dacheux, D., Albisetti, A., Byard, E.H., Bonhivers, M., and Robinson, D.R. (2015). BILBO1 Is a Scaffold Protein of the Flagellar Pocket Collar in the Pathogen Trypanosoma brucei. PLoS pathogens 11, e1004654.

75. Selvapandiyan, A., Duncan, R., Debrabant, A., Bertholet, S., Sreenivas, G., Negi, N.S., Salotra, P., and Nakhasi, H.L. (2001). Expression of a mutant form of Leishmania donovani centrin reduces the growth of the parasite. J Biol Chem 276, 43253–43261.

76. Absalon, S., Blisnick, T., Kohl, L., Toutirais, G., Dore, G., Julkowska, D., Tavenet, A., and Bastin, P. (2008). Intraflagellar transport and functional analysis of genes required for flagellum formation in trypanosomes. Mol Biol Cell 19, 929–944.

77. Kohl, L., Sherwin, T., and Gull, K. (1999). Assembly of the Paraflagellar Rod and the Flagellum Attachment Zone Complex During the Trypanosoma brucei Cell Cycle. Journal of Eukaryotic Microbiology 46, 105–109.

78. Devaux, S., Kelly, S., Lecordier, L., Wickstead, B., Perez-Morga, D., Pays, E., Vanhamme, L., and Gull, K. (2007). Diversification of function by different isoforms of conventionally shared RNA polymerase subunits. Mol Biol Cell 18, 1293–1301.

79. Hombach, A., Ommen, G., Chrobak, M., and Clos, J. (2013). The Hsp90–Sti1 interaction is critical for Leishmania donovani proliferation in both life cycle stages. Cellular Microbiology 15, 585–600.

80. Schindelin, J., Arganda-Carreras, I., Frise, E., Kaynig, V., Longair, M., Pietzsch, T., Preibisch, S., Rueden, C., Saalfeld, S., Schmid, B., et al. (2012). Fiji: an open-source platform for biological-image analysis. Nature methods 9, 676–682.

81. de Chaumont, F., Dallongeville, S., Chenouard, N., Herve, N., Pop, S., Provoost, T., Meas-Yedid, V., Pankajakshan, P., Lecomte, T., Le Montagner, Y., et al. (2012). Icy: an open bioimage informatics platform for extended reproducible research. Nature methods 9, 690–696.

82. Dufour, A., Meas-Yedid, V., Grassart, A., and Olivo-Marin, J.- (2008). Automated quantification of cell endocytosis using active contours and wavelets. 19th International Conference on Pattern Recognition, 1-4.

83. Dufour, A., Thibeaux, R., Labruyere, E., Guillen, N., and Olivo-Marin, J.C. (2011). 3-D active meshes: fast discrete deformable models for cell tracking in 3-D time-lapse microscopy. IEEE transactions on image processing: a publication of the IEEE Signal Processing Society 20, 1925–1937.

