## supplemental figure S1 to S6 for "The low complexity regions in the C-terminus are essential for the subcellular localisation of *Leishmania* casein kinase 1 but not for its activity"

### Figure S1 – CK1.2-V5 localisation in methanol-fixed promastigotes

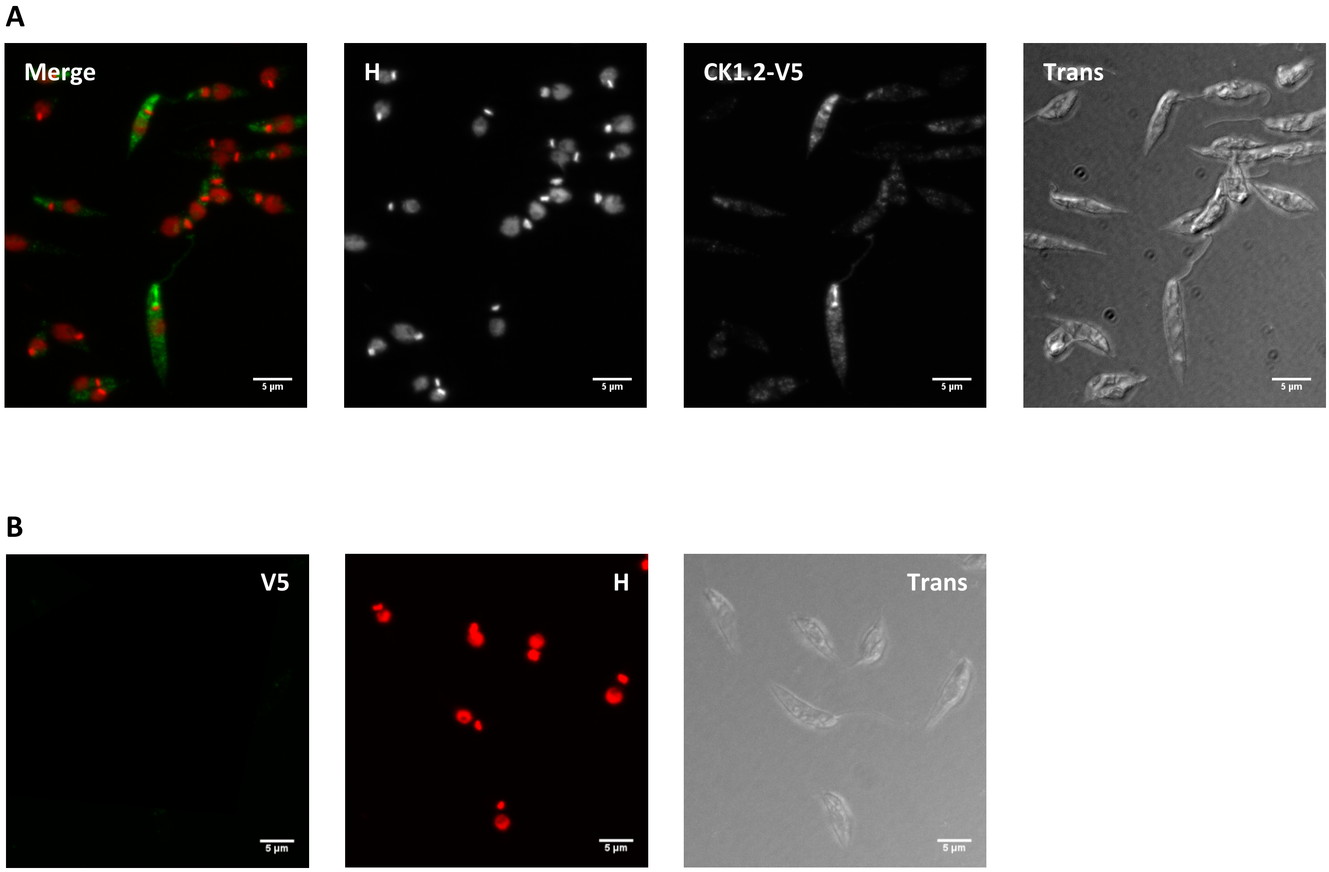

### Figure S2 – Hsp90/HSP70 cytoplasmic localisation

**A**

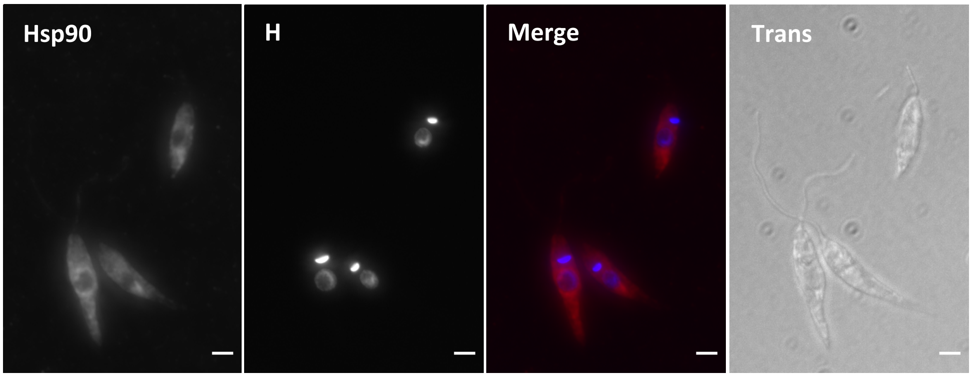


**B**

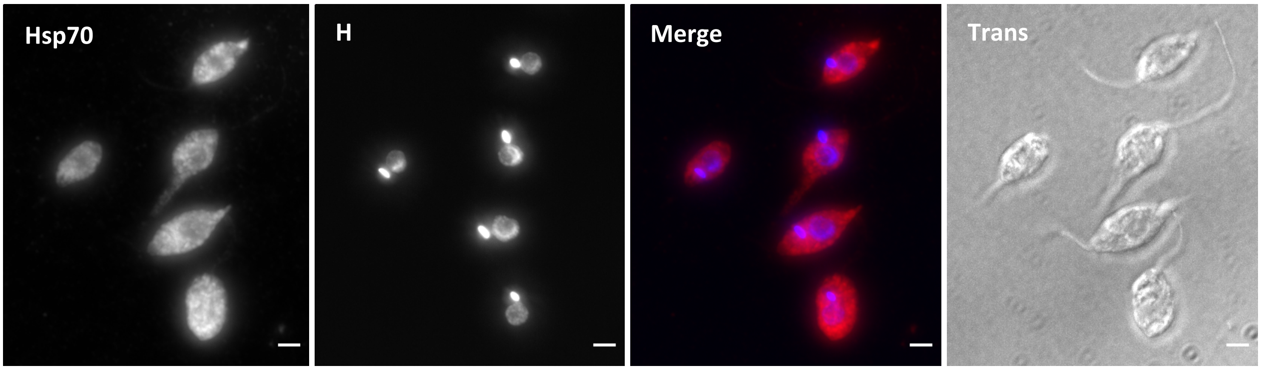

Figure S3 – LmCK1.2 is redistributed to the mitotic spindle during mitosis

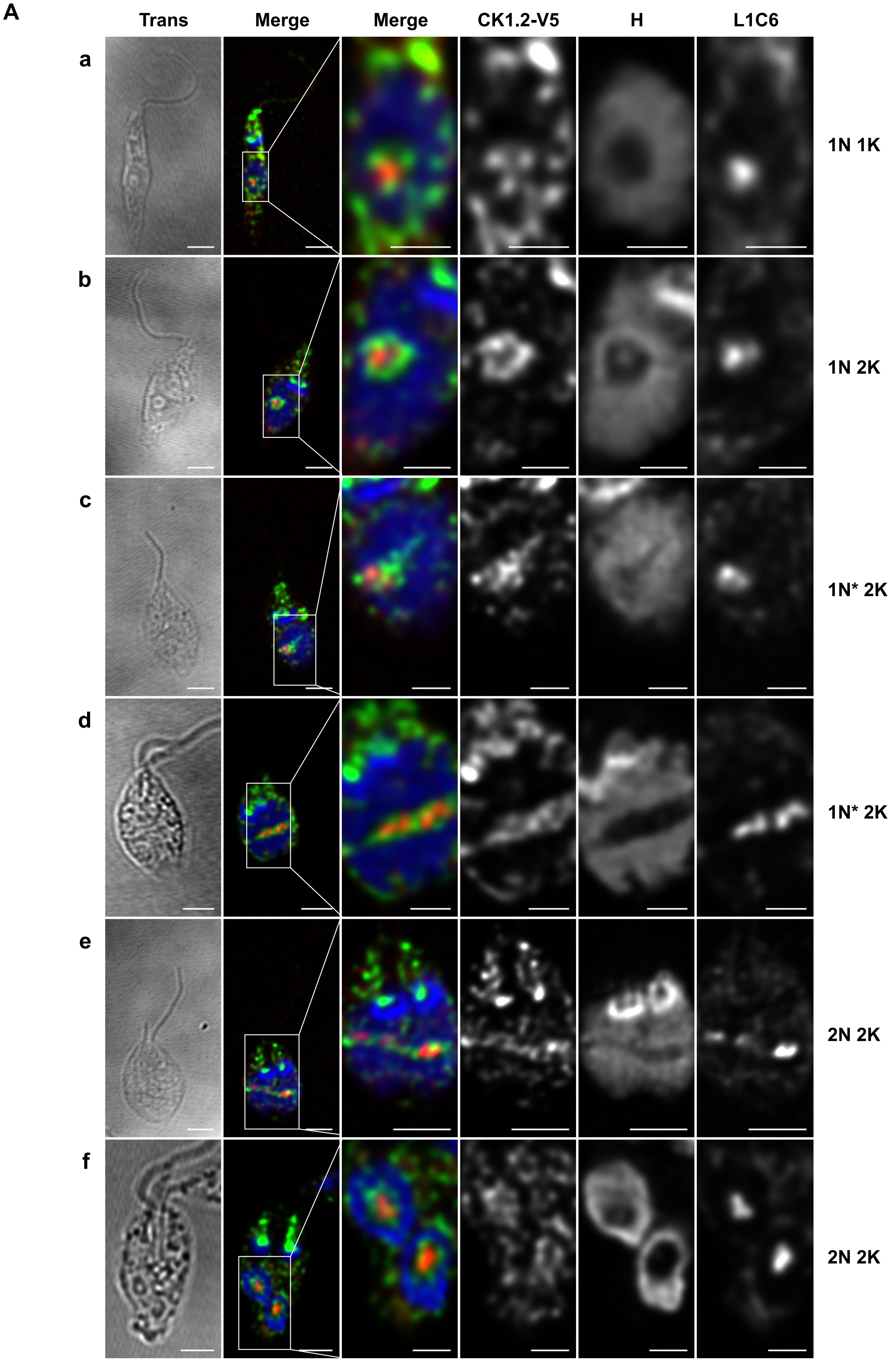

Figure S3 – LmCK1.2 is redistributed to the mitotic spindle during mitosis

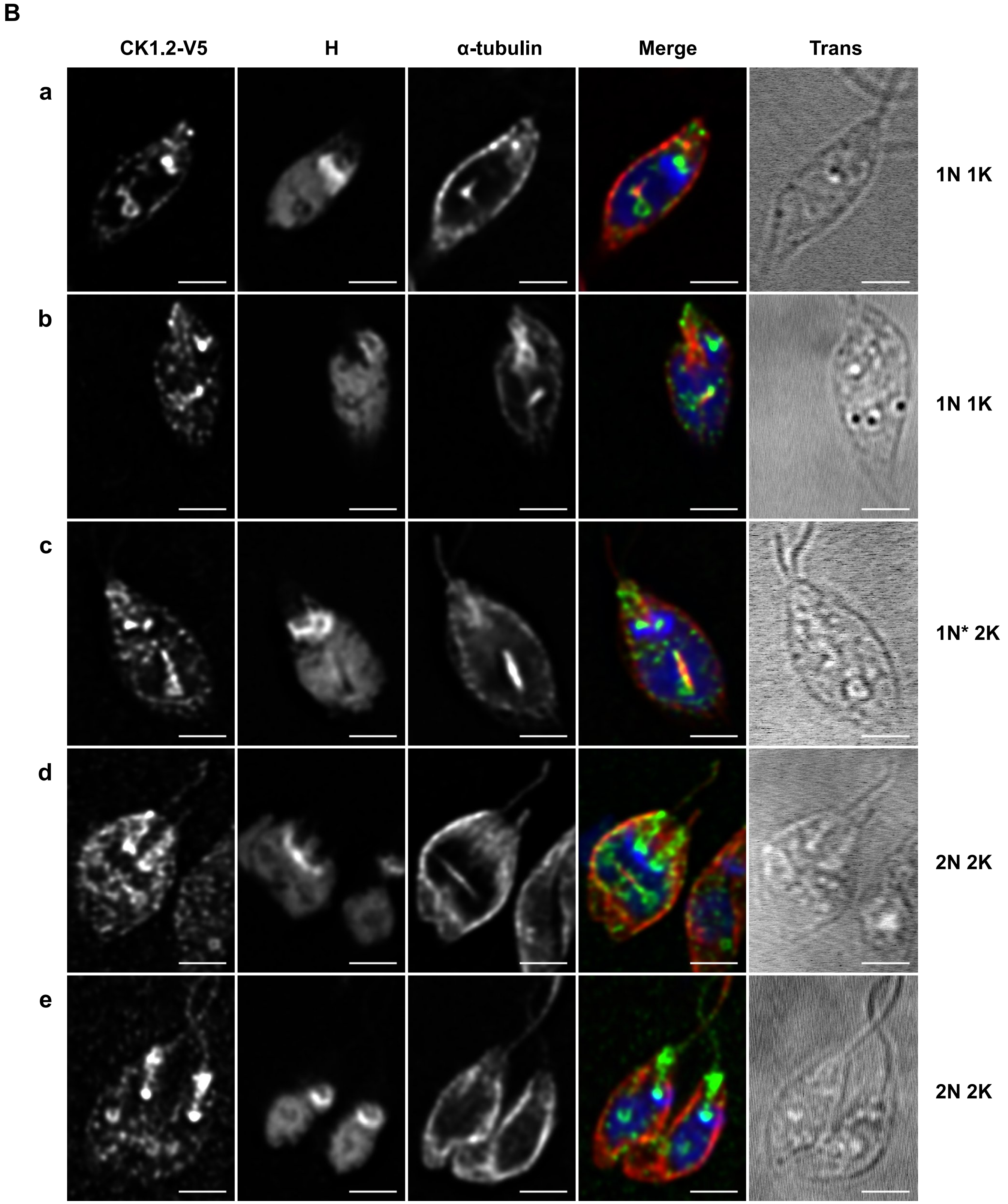

### Figure S4 – Inhibition of proteasomal and lysosomal degradation

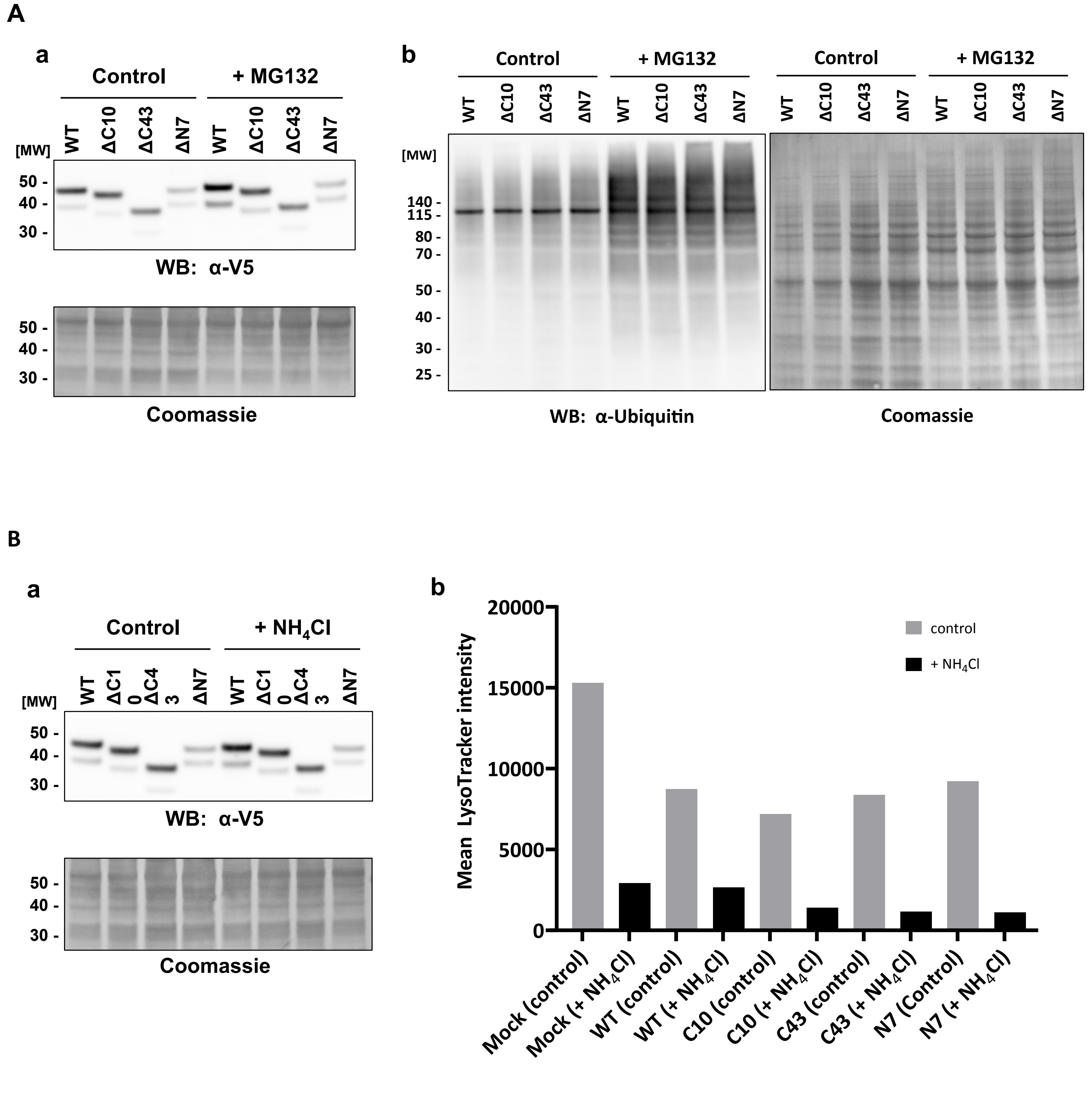

Figure S5 – Position of the LCR in human CK1 othologs.

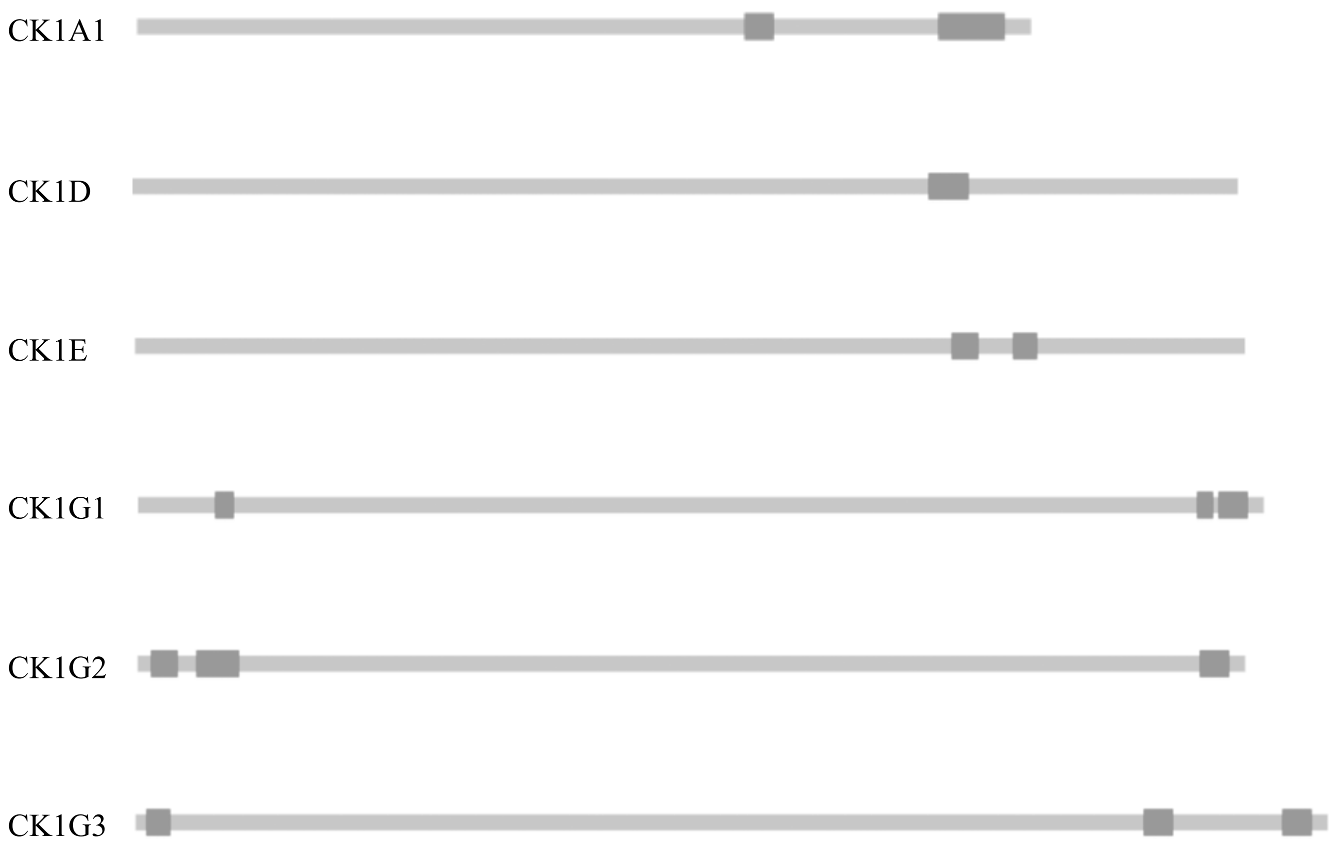

### Figure S6 – Protocol for automatic segmentation of parasite bodies.

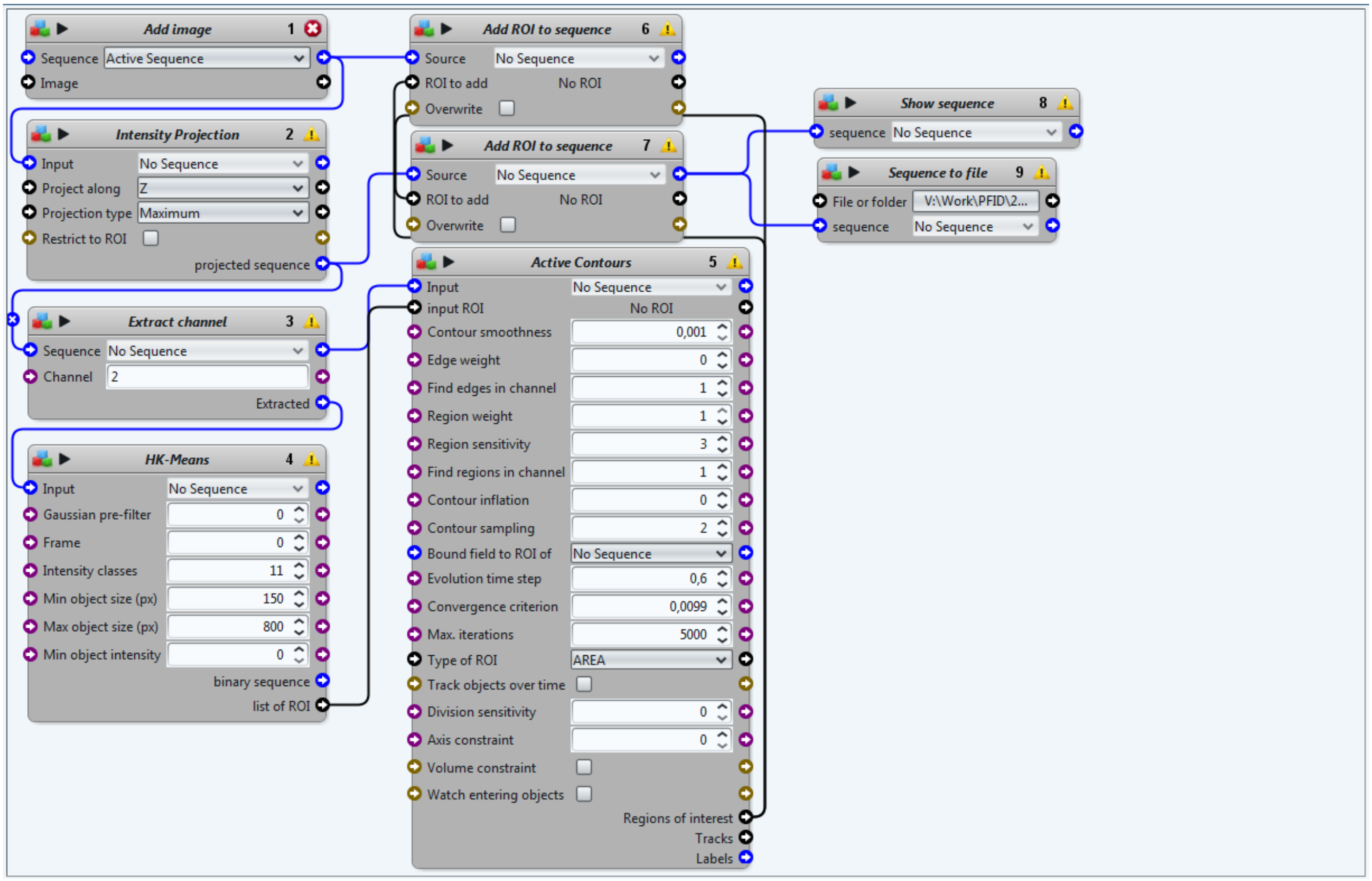
